# Kinetic Implications of IP_6_ Anion Binding on the Molecular Switch of the HIV-1 Capsid Assembly

**DOI:** 10.1101/2024.12.05.627050

**Authors:** Manish Gupta, Arpa Hudait, Mark Yeager, Gregory A. Voth

**Author notes:** equal contribution (Manish Gupta, Arpa Hudait).

## Abstract

HIV-1 capsid proteins (CA) self-assemble into a fullerene-shaped capsid, enabling cellular transport and nuclear entry of the viral genome. A structural switch comprising the Thr-Val-Gly-Gly (TVGG) motif either assumes a disordered coil or a 3_10_ helix conformation to regulate hexamer or pentamer assembly, respectively. The cellular polyanion inositol hexakisphosphate (IP6) binds to a positively charged pore of CA capsomers rich in arginine and lysine residues mediated by electrostatic interactions. Both IP6 binding and TVGG coil-to-helix transition are essential for pentamer formation. However, the connection between IP6 binding and TVGG conformational switch remains unclear. Using extensive atomistic simulations, we show that IP6 imparts structural order at the central ring, which results in multiple kinetically controlled events leading to the coil- to-helix conformational change of the TVGG motif. IP6 facilitates the helix-to-coil transition by allowing the formation of intermediate conformations. Our results identify the key kinetic role of IP6 in HIV-1 pentamer formation.

## Introduction

HIV-1 capsid maturation involves sequential cleavage of the Gag polyprotein, leading to a conical capsid assembly with the liberated CA protein ^1-4^. The mature capsid protects the viral RNA from the cytosolic immune defense during transport within the cell and mediates nuclear import through the nuclear pore complex (NPC) ^5-9^. The conical shape of the capsid is particularly relevant for nuclear entry, reverse transcription of the viral genome in the nucleus, and release of the DNA at the sites of integration ^9-11^. The HIV-1 cone can dock at the NPC central channel when approaching from the narrow end, which allows the pore to progressively dilate to relieve stress as the scaffold nucleoporins encounter the wider sections of the capsid ^6,8^. HIV-1 capsid utilizes varying surface curvature to selectively interact with cellular host factors ^12,13^. When the conical capsid is docked at the NPC central channel, the host factor NUP153 prevalent at the nuclear basket interacts with the narrow end to facilitate translocation into the nucleus. These features suggest that there is a distinct evolutionary advantage of the conical shape of the capsid for productive viral infection.

The conical capsid comprises hexamers with exactly 12 pentamers at the wide and narrow ends to stabilize the curvature required for capsid closure (13, 14). The CA monomers consist of the N-terminal domain (CA_NTD_) and C-terminal domain (CA_CTD_). CA_NTD_ consists of seven *α*-helices and a *β*-hairpin, the CA_CTD_ consists of four α-helices. The CA_NTD_ and CA_CTD_ are connected by a linker, which imparts conformational flexibility ^14^. The CA monomer leverages conformational flexibility to control quasi-equivalent capsomer association, imparting varying curvature in the capsid lattice^14-16^. The interactions between CA_NTD_ allow the formation of the capsomers, while the interactions between CA_CTD_ regulate the assembly of hexamers and pentamers to form the capsid lattice. The small molecule polyanionic IP_6_ is an assembly cofactor for both immature and mature HIV-1 CA lattices ^17-20^. Structural characterization of the mature capsid identified that IP_6_ coordinates R18 residues in the central pore of CA_NTD_, enabling nucleotide diffusion into the capsid core for reverse transcription ^21^. Additionally, IP_6_ can also bind to the ring of K25 residues below the R18 ring ^22^. During in vitro reconstitutions of CA assembly, the addition of IP_6_ promotes cone-shaped morphologies by coordinating the positively charged ring of R18 and K25 in the central pore of both hexamer and pentamers ^17,18,23^. In the pentamers, IP_6_ can bind to R18 and K25 rings with comparable probability. In contrast, in the hexamers, IP_6_ preferentially binds to the R18 ring since the pore size for the K25 ring is significantly larger ^12^. Recent cryo-electron microscopy (cryo-EM) and molecular dynamics simulations demonstrated that pentamer formation is significantly dependent on the reduction of electropositive repulsion in the central pore by IP_6_ binding ^23-25^. Free energy quantification reveals a more stable binding of IP_6_ to R18 in CA pentamers relative to hexamers ^23,25^. The stabilization of IP_6_ is facilitated by smaller pore volume in pentamer, which allows tighter coordination ^25^.

The relative arrangement of CA_NTD_ and CA_CTD_ is mostly identical in hexamers and pentamers. However, there is a significant difference between the conformation of the base of helices 3 and 4 and the loop connecting the helices. The TVGG motif (residues 58 to 61) at the base of helix 3 is in coil conformation in hexamer. In contrast, that motif forms a 3_10_ helix in the pentamer ^12,26^. The TVGG motif is juxtaposed at the NTD-NTD and NTD-CTD interfaces that maintain capsomer stability ^27,28^. Interprotomer NTD-NTD interface involves contact between P38/M39 in helix 2 and N57/T58 in helix 3 of neighboring subunits in the hexamer ^27^, whereas P38/M39 contact V24/K25 in the adjacent helix 1 in the pentamer ^26^. Notably, cryo-EM and biochemical analyses revealed that a structural switch in the TVGG motif of CA modulates the relative distribution of hexamers and pentamers in the polyhedral capsid ^26^. Furthermore, the 3_10_ helix formation results in NTD-NTD and NTD-CTD interactions that are unique to CA pentamers.

The structural implications of IP_6_ binding to the pentamer and TVGG coil-to-helix transition have not been known. It has been proposed that IP_6_-induced conformational switching may occur through allosteric communication through the hydrophobic core mediated by a network of interactions. Ligand-induced allostery can occur through changes in the protein structure or dynamics ^29,30^. Key questions are: Does IP_6_ binding trigger the conformational switching of the TVGG motif? If IP_6_ influences the conformational switching, what structural and dynamic alternation of the CA subunit does it induce that culminates in the coil-to-helix transition?

To answer these mechanistic questions, we performed extensive unbiased all-atom (AA) molecular dynamics (MD) simulations, conformational sampling, and well-tempered metadynamics (WT-MetaD) ^31-33^ biased free energy sampling simulations to investigate the network events triggered by the binding of IP_6_, and conformational switching of the TVGG motif. In the absence of IP_6_, the R18 residues remain disordered to elude electrostatic repulsion in the pore. When IP_6_ occupies the pore of the pentamer, it is localized at a spot midway between the plane of the R18 and K25 rings, coordinating positively charged moieties of the R18 and K25 residues. In this arrangement, the R18 residues are pointed downwards, and the K25 residues point upwards. In the presence of IP_6_, the K25 residues are conformationally rigidified, allowing stable K25-M39 contacts. Thus, in the presence of IP_6_, M39 points away from the TVGG motif, which creates a pocket at the base of helix 3. The formation of this pocket provides requisite room for the TVGG motif in a coil conformation to fold into a partial 3_10_ helix. Our results identify a three-step mechanism initiated by IP_6_ binding to CA pentamers that results in the folding of the TVGG random coil to the 3_10_ helix. Both during unbiased and WT-MetaD simulations in the presence of IP_6_, the TVGG motif undergoes the coil-to-helix transition mediated via a partially folded intermediate state. Our results uncover several key mechanistic details of how IP_6_ regulates TVGG motif conformational switching in CA pentamers through allostery. These results also may point to new druggable targets that could impede HIV-1 capsid formation.

## Results

### Molecular mechanism of TVGG motif coil-to-helix transition in CA pentamers

Our atomic model of the mature HIV-1 capsid system contains roughly 100 million atoms ^34^, making it computationally intractable for exhaustive, unbiased AA MD simulations. Instead, to limit computational expense, we simulated a global pentamer structure to investigate the relationship between IP_6_ binding and the TVGG coil-to-helix transition. Here, the “global pentamer” patch comprised a central pentamer surrounded by five peripheral hexamers (Fig. 1A) in the solvent. To simulate the IP_6_ effect, another global pentamer system was prepared by adding IP_6_ anions roughly 3 Å above the central R18 ring in each capsomere. In total, four replicas of the global pentamer were simulated for a total of ∼12 μs with and without IP_6_.

**Fig 1.**
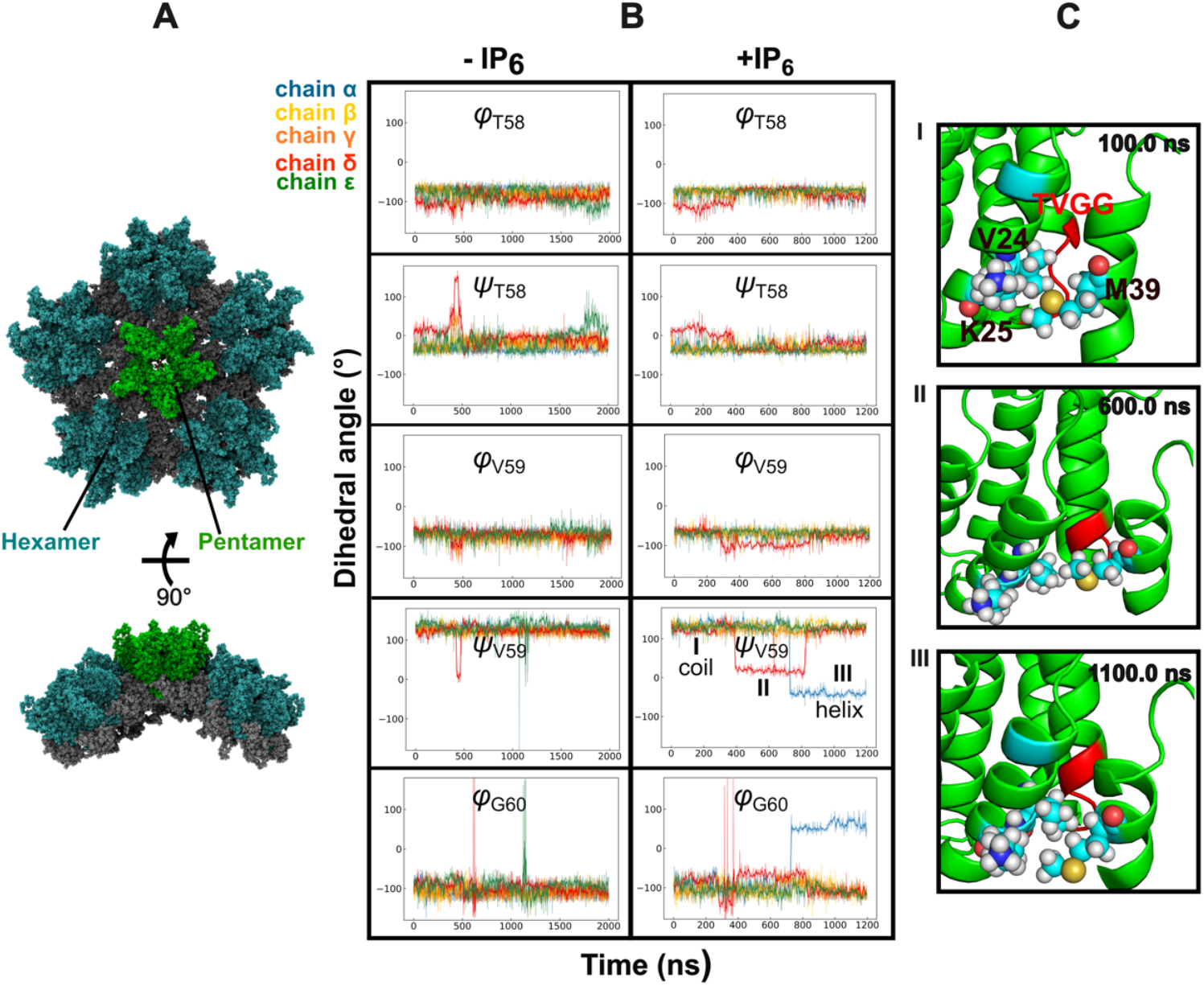
IP_6_ facilitates coil-to-helix transition in the TVGG motif of the CA pentamer. **(A)** The **‘**global pentamer’, made of a central CA pentamer (green) and five adjacent hexamers (blue), forms a curved lattice. **(B)** Time series of TVGG backbone (φ, ψ) dihedral angles of each chain in the central pentamer with (left panel) and without IP_6_ (right panel) for times longer than 100 ns. In the absence of IP_6_, the TVGG (φ, ψ) dihedral angle space briefly explored a metastable state but primarily stayed in the coil conformation. In all the IP_6_-bound pentamer simulations, the TVGG motif transitioned to either to a metastable state with ψ_*V59*_ and φ_*G60*_ values of ∼10° and -60° respectively or to a partial helix conformation with ψ_*V59*_ and φ_*G60*_ values of -45° and 60° respectively. **(C)** Snapshots from IP_6_-bound global pentamer MD trajectory show the conformational transition in the TVGG motif of the CA pentamer. Zoomed-in view of the TVGG motif (red) of α-subunit in the coil conformation is shown in the panel-I at the top. During the course of the simulation, TVGG motif in the δ-subunit and α-subunit in the pentamer transition to a metastable (II) and partial 3_10_ helix (III) conformation, respectively. Timesteps at which the snapshots are recorded are labeled at the upper-right corner of each panel.

We systematically analyzed the structural changes in the TVGG motif of the central pentamer of the capsid patch. In the initial pentamer structure, the TVGG motif resembled a random coil configuration, equivalent to the conformation in hexamers. Although the disordered conformation of the TVGG motif is termed “random coil”, there is an ensemble of backbone geometries in the (φ, ψ) dihedral angle space (Fig. 1B). The pentamer TVGG backbone (φ, ψ) dihedral angles fluctuated persistently in the absence of IP_6_. T58 (φ, ψ) dihedral angles gradually converged to approximate values of - 60° and - 45°, respectively, which are indicative of a 3_10_ helix conformation. Notably, φ_*V*59_ resembled a helix-like structure with a value of -60°, but ψ_*V*59_ value of ∼125° deviated from 3_10_ helix. φ_*G60*_ converged to a large negative value of -100°, but ψ_*G60*_ and G61 (φ, ψ) adopted a random coil conformation.

In the presence of IP_6_, the fluctuation of the TVGG backbone (φ, ψ) dihedral angles are significantly altered. T58 (φ, ψ) and φ_*V*59_ dihedral angles resembled a 3_10_ helix. However, major transitions in ψ_*V*59_ and φ_*G60*_ values were observed in the α-domain. ψ_*V*59_ and φ_*G60*_ transitioned to a value of - 45° and 60° (Fig. 1B) respectively to allow a coil-to-helix switch within the TVGG motif (Fig. 1C). In the δ-domain, ψ_*V*59_ transitioned to a value of 10°, and φ_*G60*_ concurrently transitioned to -60°, indicating the existence of a transiently populated and partially folded intermediate state between the coil and 3_10_ helix. The relaxation of ψ_*V*59_ to a 3_10_ helix configuration was more pronounced for pentamers with IP_6_, suggesting that the TVGG coil-to-helix switch might be directly related to IP_6_ binding at the central channel. The anionic phosphate groups of IP_6_ remain bound to the R18 ring throughout the duration of the simulation. In all except one IP_6_-bound pentamer simulations, ψ_*V*59_ switched to a value of -45°, and φ_*G60*_ switched to 60° concomitantly, assuming a 3_10_ helix structure.

### IP_6_ imparts order in the positively charged R18 and K25 rings

The CA hexamer and pentamers contain two electropositive rings composed of R18 and K25 residues, mutually stacked in the central pore. The abundance of basic residues renders the pore strongly electropositive inducing asymmetry at the center ^13^. Visual inspection of the simulation trajectories revealed that R18 residues are highly orientationally dynamic (Fig. 2A). To characterize the orientational fluctuations, we estimated the variation in root-mean-square fluctuation (RMSF) of the helix-1 center of mass relative to the other helix-1 in the pore (Fig. 3C). RMSF calculations revealed a persistent disorder in the central ring of the pentamer in the absence of IP_6_. Evidently, the R18 chain fluctuated between two conformations, directing either towards the exterior or interior of the capsid. Snapshots of the pentamer central pore with and without IP_6_ are shown in Fig. 2A and 2B, respectively, and the dynamics are shown in movies S1 and S2. The orientational disorder of the R18 residues is instigated by the electrostatic repulsion between the charged guanidino groups at the pore. Consequently, the guanidino groups fluctuate rapidly to orient in the opposite direction of the adjacent positively charged moieties of arginine.

**Fig 2.**
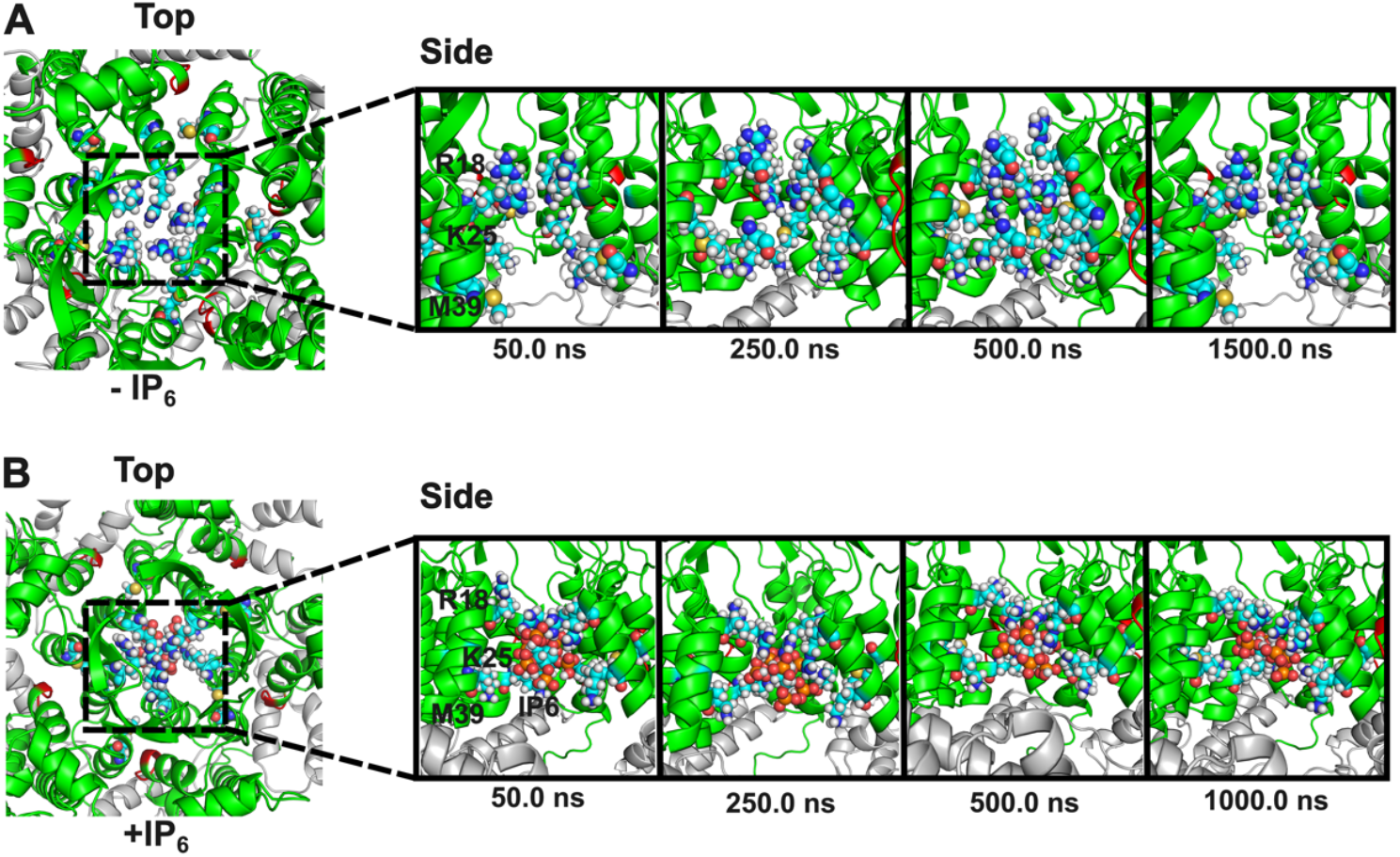
Comparison of central pores from CA pentamer and IP_6_-bound pentamer. **(A)** Snapshots from (−IP_6_) simulations show that the ‘molecular iris’ made of β-hairpins is rendered asymmetric by the repulsion of positively charged residues. Close-up views of the central channel show that in the absence of any stabilizing force, R18 residues fluctuate between two different configurations, whereas K25 is oriented towards the interior of the capsid. ε-CA domain not shown for clarity. **(B)** IP_6_ imparts fivefold symmetry in the central channel. IP_6_ is bound below the R18 ring, coordinating multiple R18 and K25 residues. R18 side chains point towards the interior, and K25 side chains point towards the center of the capsid to contact IP_6_.

**Fig 3.**
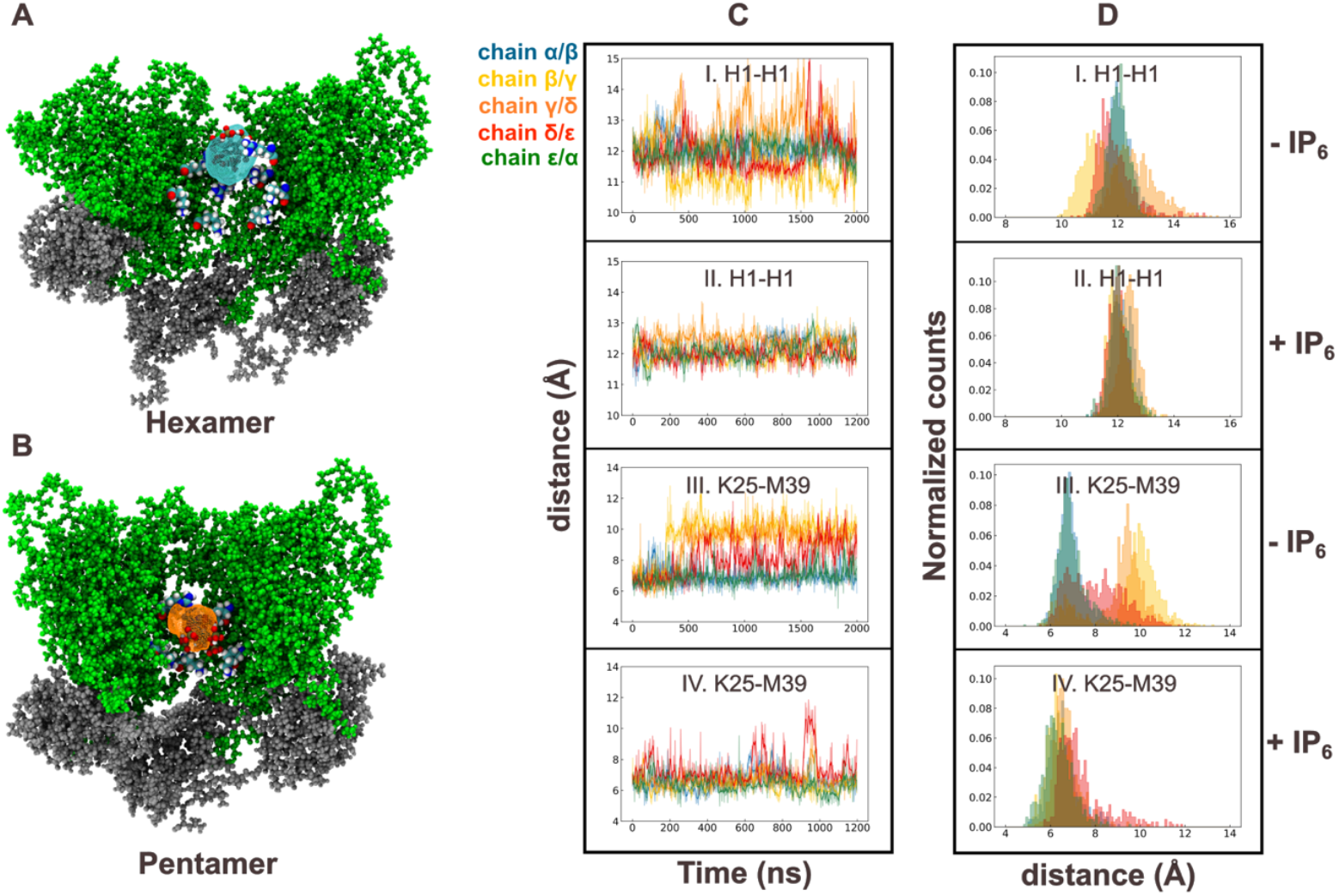
Characterization of IP_6_ binding modes and structural changes in the pentamer. **(A)** IP_6_ density map is overlayed with CA hexamer structure. Densities contribute to IP_6_ binding above R18 ring in hexamer with only a small ligand density below the R18 ring. **(B)** The ligand density map in the pentamer contains IP_6_ primarily below the R18 ring. **(C)** Time series plot of distances I. H1(*CA*_*i*_)-H1(*CA*_*i+1*_) in the pentamer, II. H1(*CA*_*i*_)-H1(*CA*_*i+1*_) in the IP_6_-bound pentamer, III. K25(*CA*_*i*_)-M39(*CA*_*i+1*_) in the pentamer, IV. K25(*CA*_*i*_)-M39(*CA*_*i+1*_) in the IP_6_-bound pentamer. **(D)** Histograms of the distances between I. H1(*CA*_*i*_)-H1(*CA*_*i+1*_) in the pentamer, II. H1(*CA*_*i*_)-H1(*CA*_*i+1*_) in the IP_6_-bound pentamer, III. K25(*CA*_*i*_)-M39(*CA*_*i+1*_) in the pentamer, IV. K25(*CA*_*i*_)-M39(*CA*_*i+1*_) in the IP_6_-bound pentamer.

In the presence of IP_6_, the guanidinium side chains of R18 were ordered and flipped toward the channel’s center (Fig. 2B). Despite the smaller pentamer pore size, IP_6_ could not stabilize all five R18 side chains simultaneously, leaving at least one R18 residue uncoordinated. K25 side chains were also oriented towards the center of the pore to coordinate IP_6_. Notably, IP_6_ binding had a significant influence on the central pore, as its dynamic fluctuations were reduced (Fig. 3C). Previous ligand density maps have identified the metastable interactions that guide free IP_6_ toward the capsid central pore. However, longer simulations of IP_6_ bound capsid were required to access accurate spatial distribution and dynamics of IP_6_ in the binding pocket. We calculated a 3D ligand density map and compared the IP_6_ binding modes at the hexamer and pentamer central pore. We observed that IP_6_ was preferably located above the R18 ring, which is known as the primary binding site in the hexamer pore. Only a weak IP_6_ ligand density was observed below the R18 ring, likely due to the relatively larger diameter of the K25 ring for the hexamer. Intriguingly, significant IP_6_ density was observed below the R18 ring, highlighting the differential mode of binding in the pentamer central pore.

### IP_6_ binding allows M39 packing at the V24/K25 hole in pentamers

Recent high resolution cryo-EM structures have identified distinctly different packing at the NTD-NTD interface for HIV-1 hexamer and pentamer ^26^. NTD-NTD interaction between adjacent CA subunits in the hexamer is mediated by contact between M39 in helix 2 and T58 in helix 3. In CA pentamers, M39 in helix 2 preferentially contacts K25 in helix 1 ^35^. In our unbiased AA MD simulation trajectories, we analyzed the molecular contacts of K25, M39, and T58 in the presence or absence of IP_6_.

The time series plots of the K25(*CA*_*i*_)-M39(*CA*_*i+1*_) and M39(*CA*_*i*_)-T58(*CA*_*i+1*_) distances between adjacent subunits are shown in Fig. 3B and Fig. 4A, respectively. In the absence of IP_6_, the RMS distance between the center of mass of K25-M39 between adjacent CA subunits increases. Simultaneously, the M39-T58 distances also decrease, which indicates the formation of contact. The distribution of the K25-M39 fluctuations revealed that K25-M39 packing is not favored in the absence of IP_6_ (Fig. 3C). In the absence of IP_6_, the disordered central pore prevented the stable packing of the M39 residue in the hole formed by V24 and K24, even though this configuration is preferred in pentamers. Histograms of H1(*CA*_*i*_)-H1(*CA*_*i+1*_) and K25(*CA*_*i*_)- M39(*CA*_*i+1*_) distances in Fig. 3D distinctly demonstrate that the fluctuations within the H1-H1 contacts in the β*/γ*, γ/δ, and δ*/*ε CA domains are relayed to K25-M39 contacts in the same subunits. The loss of K25-M39 contact moved the M39 residue toward the N57 and T58, forming a hexamer-like NTD-NTD interface (Fig. 4A and 4C).

**Fig 4.**
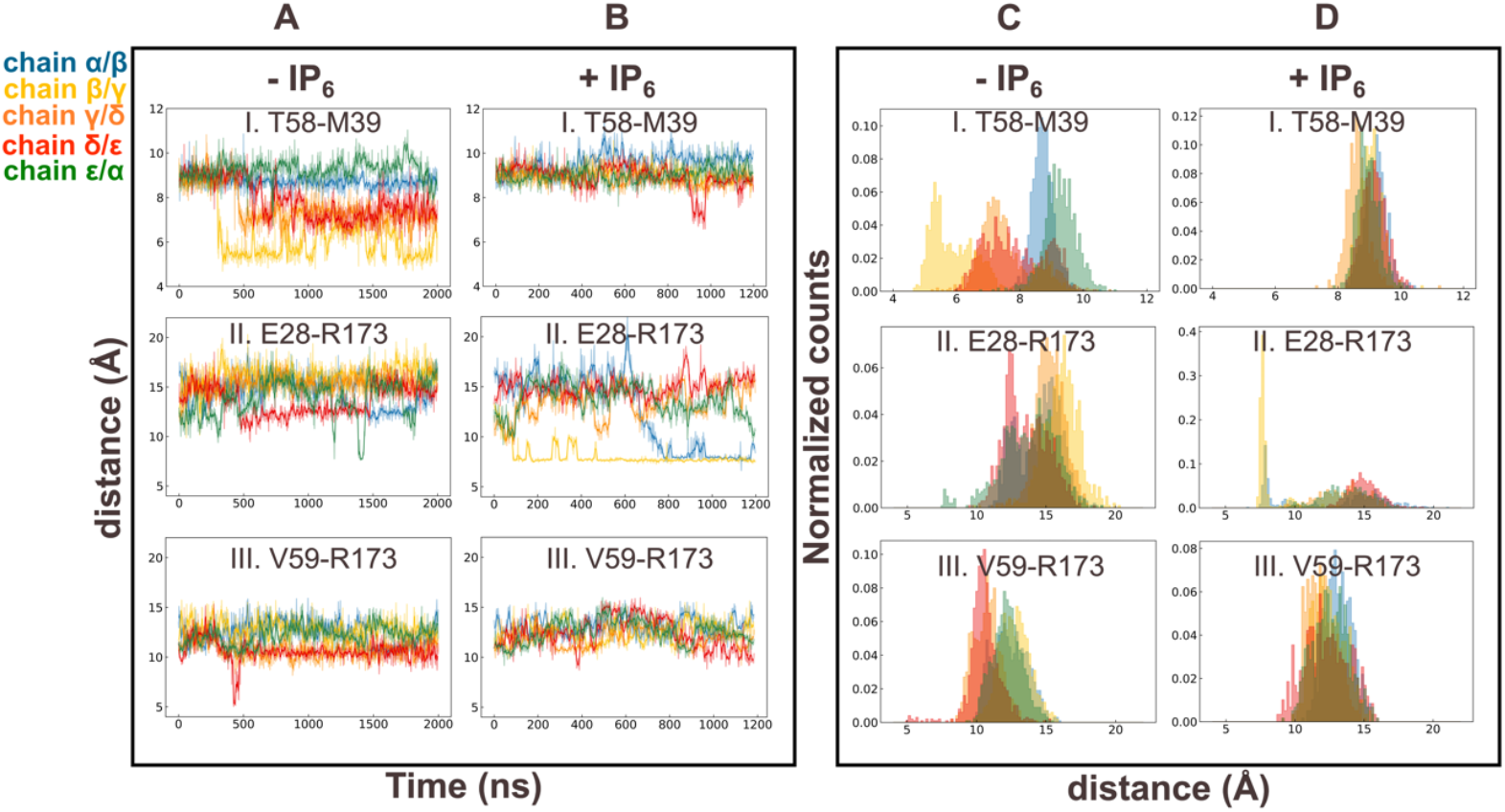
Structural changes in NTD-NTD and NTD-CTD interfaces in the CA pentamer. **(A)** Time courses of I. T58(*CA*_*i*_)-M39(*CA*_*i+1*_) interaction and II. E28(*CA*_*i*_)-R173(*CA*_*i+1*_) interaction and III. V59(*CA*_*i*_)-R173(*CA*_*i+1*_) interaction in the pentamer. **(B)** Time courses of I. T58(*CA*_*i*_)-M39(*CA*_*i+1*_) interaction and II. E28(*CA*_*i*_)-R173(*CA*_*i+1*_) interaction and III. V59(*CA*_*i*_)-R173(*CA*_*i+1*_) interaction in the IP_6_-bound pentamer. **(C)** histogram of distances between I. T58(*CA*_*i*_)-M39(*CA*_*i+1*_) and II. E28(*CA*_*i*_)-R173(*CA*_*i+1*_) and III. V59(*CA*_*i*_)-R173(*CA*_*i+1*_) in the pentamer. **(D)** histogram of distances between I. T58(*CA*_*i*_)-M39(*CA*_*i+1*_) and II. E28(*CA*_*i*_)-R173(*CA*_*i+1*_) and III. V59(*CA*_*i*_)-R173(*CA*_*i+1*_) in the IP_6_-bound pentamer.

To summarize, in the absence of IP_6_, the molecular environment in the pentamer drifts towards the hexamer. Fluctuations of residues adjacent to the central pore of pentamers are significantly reduced in the presence of IP_6_ (Fig. 3C). This allowed stable packing of M39 against the V24/M39 hole, trapping it in the pentamer state. The formation of molecular contacts between helix 1 and 2 excludes helix 3 from the NTD-NTD packing, creating an empty pocket and allowing the TVGG motif to fold into a 3_10_ helix. A similar trend is observed in all four replica simulations (Fig. S1-S12 of Supporting Information).

The TVGG motif is located at the NTD-CTD interface that stabilizes the intra-capsomer contacts between adjacent subunits. The pocket is the binding site for host cell factors responsible for nuclear import and small molecule inhibitors that target the HIV-1 capsid ^36,37^. In the hexamer, V59 of the TVGG motif interacts with R173 of the neighboring CA domain, contributing to the packing of helix 3 and helix 8 at the NTD-CTD interface ^12,26^. In contrast, structural changes in the pentamer trigger loss of contact between V59(*CA*_*i*_)-R173(*CA*_*i+1*_), allowing R173 to engage with E28 in the adjacent helix 1. In the capsid patch simulations, the distance between E28 and R173 residues in adjacent CA chains increased to a value of ∼12 Å, and the V59-R173 distance decreased in the absence of IP_6_, signifying a tendency to adopt hexameric arrangement. IP_6_-bound CA pentamer, however, favored pentamer configuration with stable interprotomer helix 1-helix 8 interaction between E28 and R173 residues.

To further investigate the structural rearrangements effected by IP_6_, we compared contacts made between *CA*_*i*_ helix 1 or helix 3 and helix 2 of *CA*_*i+1*_(Fig. S13). In IP_6_-bound pentamer, residues located at the base of helix 1 engage residues 34-36 in the helix 2 of the neighboring protomer favoring pentamer specific contacts. In contrast, a hexamer-like NTD/NTD interface was adopted in the absence of IP_6_ as residues 57-60 located at the base of helix 3 slipped towards the helix 2 of adjacent CA subunit. These findings suggest that IP_6_ binding can influence the TVGG switch in the pentamer by modulating interactions within the NTD-NTD and NTD-CTD interfaces.

### Partially folded intermediates facilitate coil-to-helix transition in the presence of IP_**6**_

In long unbiased MD simulations in the presence of IP_6_, the TVGG motif fluctuates between these states: coil *→* partially folded *→* 3_10_ helix. In contrast, in the absence of IP_6_, the TVGG motif fluctuates between these states: coil *⇌* 3_10_ helix. To assist in the interpretation of the above unbiased atomistic MD simulations, we performed WT-MetaD simulations of a CA pentamer in the presence and absence of IP_6_. Briefly, WT-MetaD simulation is an enhanced sampling method that can accelerate the timescales of transitions between states, which are often not sampled in unbiased simulations ^32,38^. WT-MetaD simulations are performed as a function of appropriate reaction coordinates (often called “collective variables”, or CVs), which provide an energetic landscape of the conformational transition. In the WT-MetaD simulations, the central pentamer is surrounded by partial hexamers (see Methods for details of system construction). The reaction coordinates in the WT-MetaD simulations are the distances between K25 of *CA*_1_ and M39 of *CA*_2_ denoted as *d*_*K25-M39*_, and the helicity parameter (*h*_*V*59_) calculated from ψ_*V*59_ of *CA*_1_ (time series shown in Figs. S14 and S16). The helicity 0 and 1 corresponds to the coil and helix conformations. The choice of the reaction coordinates is informed by the analysis of the unbiased simulations. The potential of mean force (PMF, i.e., conditional free energy) of TVGG motif folding (Fig. 5A and Fig. S15; energy scale 0.2 kcal/mol) is presented as a function of the reaction coordinates *h*_*V*59_ and *d*_*K25-M39*_.

**Fig 5.**
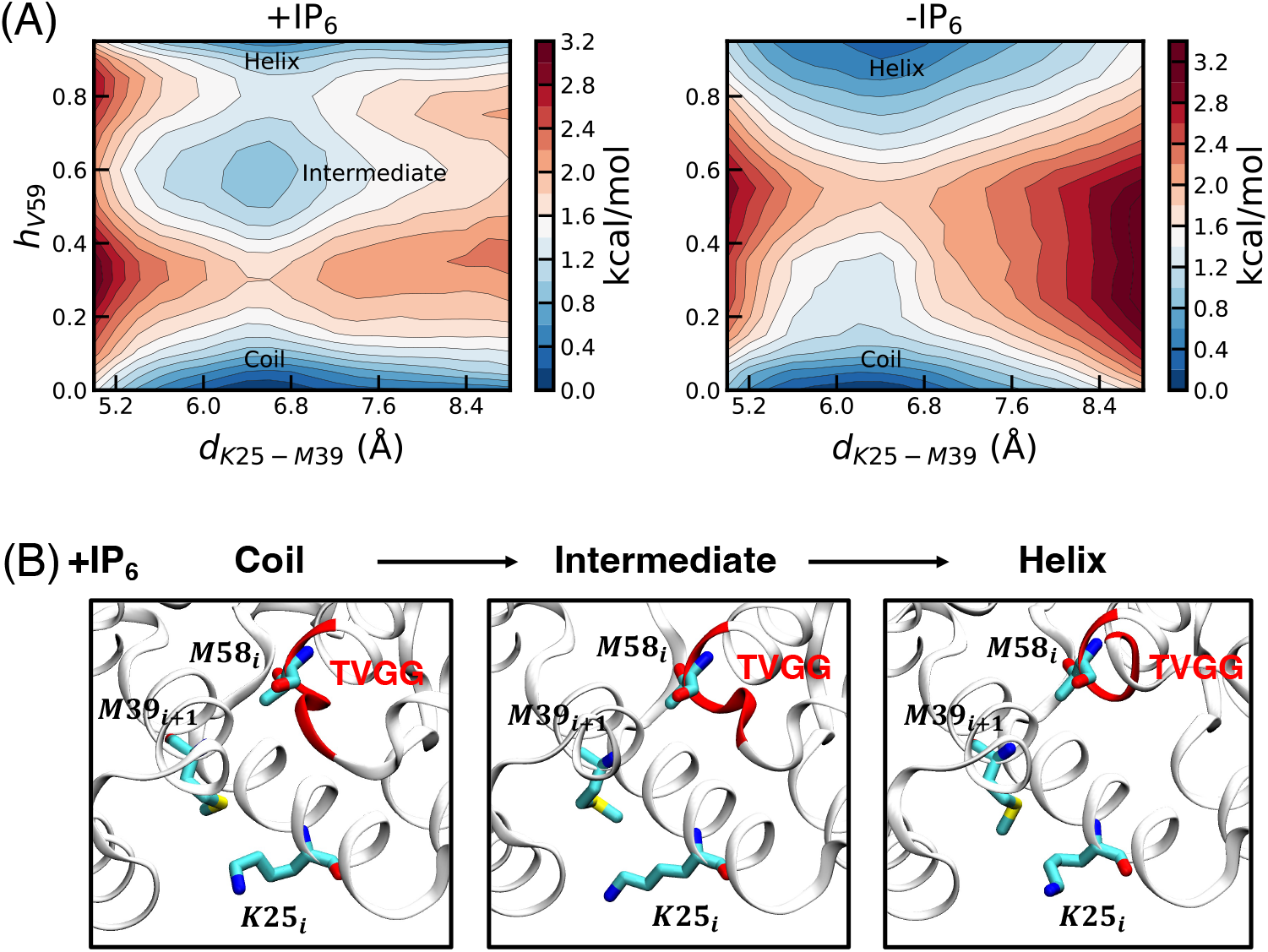
Molecular pathway of coil-to-helix transition in the CA pentamer modulated by IP_6_. **(A)** 2-dimensional plot of the PMF of the conformation landscape of the TVGG motif as a function of the helicity (*h*_*V59*_) of V59, and distance between K25 of *CA*_1_ and M39 of *CA*_2_ denoted (*d*_*K25-M39*_). The bar on the right has the color code for energy level contours in kcal/mole. The PMFs were calculated from the WT-MetaD simulations performed in the presence (+IP_6_) and absence of (−IP_6_). **(B)** Snapshots of the folding of the TVGG motif (shown in red ribbon) in the presence of IP_6_.

In the PMFs (both in the presence and absence of IP_6_), there is a deep energy minimum for the coil conformation of the TVGG motif. The minimum corresponds to a *d*_*K25-M39*_ value ranging from 6-6.8 Å. While remaining in the coil conformation, fluctuations of the K25(*CA*_1_)-M39(*CA*_2_) contact can lead to higher *d*_*K25-M39*_ values. When in the helix conformation, the energetically stable basin corresponds to *d*_*K25-M39*_ value ranging from 6-6.8 Å. Similar to that in the coil state, fluctuations allow for the K25(*CA*_1_)-M39(*CA*_2_) contacts to weaken, exploring higher *d*_*K25-M39*_ values. Therefore, both coil and helix conformations allow for heterogeneous distribution of the K25-M39 contact, consistent with the results of the unbiased simulations. Both in the presence and absence of the IP_6_, the coil-to-helix transition occurs through a funnel-like landscape. As the TVGG motif in the coil conformation begins to fold, along the reaction coordinates *h*_*V*59_, the K25(*CA*_1_)-M39(*CA*_2_) distance fluctuations narrow to the 6-6.8 Å. In the absence of IP_6_, the barrier corresponds to *h*_*V*59_ value of ∼0.5. The barrier of conformational change from coil to 3_10_ helix is ∼2 kcal/mol. As the barrier is crossed along the *h*_*V*59_ coordinate, the K25(*CA*_1_)-M39(*CA*_2_) distances gradually become more heterogeneous. In the presence of IP_6_, the barrier corresponds to *h*_*V*59_ value of ∼0.3. Interestingly, beyond this barrier, there is a distinct intermediate state corresponding to the *h*_*V*59_ value range of ∼0.52-0.65. The barrier to forming the partially folded intermediate is ∼2 kcal/mol, while forming the 3_10_ helix from the intermediate state is essentially barrierless. The presence of this partially folded state is observed in the presence of IP_6_, consistent with the observations of the unbiased MD simulations. The PMFs demonstrate that IP_6_ does not impact the energetic barrier of coil-to-helix transition. Instead, IP_6_ modulates the pathway of coil-to-helix transition via an intermediate partially folded state.

The CA monomers (except for the TVGG motif) are shown in silver ribbon. The residues *K*25_*i*_, *M*58_*i*_, and *M*39_*i+1*_ are highlighted in the snapshots.

It is informative to examine whether the intermediate partially folded conformation constitutes a metastable state in the presence of IP_6_. To this end, we picked 5 independent configurations at the intermediate state basin from the WT-MetaD simulation trajectory. We simulated these configurations for 1000 ns to examine the evolution of the intermediate-to-helix or intermediate-to-coil state populations. The system starts at *t* = 0 at the intermediate state with *h*_*V*59_ value ranging from ∼0.54-0.69. In one replica (Fig. 6), the TVGG motif continuously jumped between the helix, coil, and intermediate states. Typically, the residence time at each state is ∼10-30 ns, which corresponds to the fast interstate fluctuations and demonstrates the metastable nature of the partially folded intermediate. These fast interstate fluctuations can be interpreted as attempts to convert to either the disordered coil or folded helix conformations. These fluctuations persist up to 300 ns, after which the TVGG motif converts to the coil conformation and remains as such up to 1000 ns. In contrast, in another replica, the TVGG motif starting from the intermediate state converts to the helix within 10 ns and does not undergo any further helix-to-coil transition within 1000 ns (Fig. S16A).

**Fig 6.**
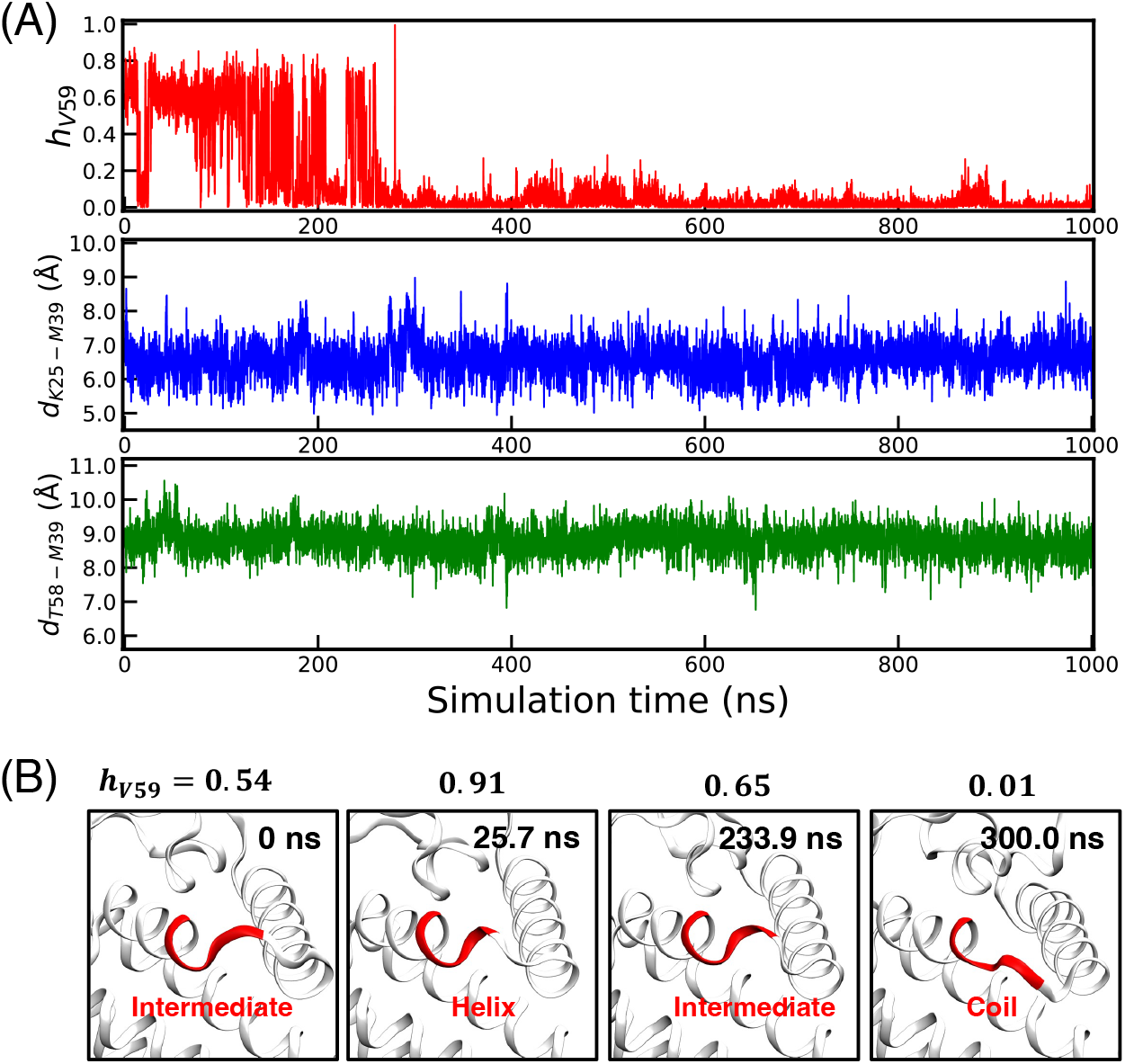
Dynamics of coil-to-helix transition from the intermediate partially folded state. **(A)** Time evolution of the helicity parameter (*h*_*V*59_), the distance between K25 and M39 (*d*_*K25-M39*_) of *CA*_1_ and *CA*_2_, and the distance between K58 and M39 (*d*_*T58-M39*_) of *CA*_1_ and *CA*_2_. The initial configuration corresponds to the intermediate partially folded state obtained from the WT-MetaD simulations in the presence of IP_6_. In the initial configuration, the values of *h*_*V*59_, *d*_*K25-M39*_, and *d*_*T58-M39*_ are 0.54, 7.2 Å, and 8.7 Å, respectively. **(B)** The snapshots show the continuous folding and unfolding of the TVGG motif (highlighted in red ribbon) within the first 300 ns simulations. The rest of the CA protein is shown in silver ribbon. For each snapshot, the title indicates the helicity parameter (*h*_*V*59_) value.

In the other three replicas, the TVGG motif at the intermediate state converts to the random coil conformation within 50 ns (Fig. S16B, S16C, and S16D). These simulations establish that the partially folded intermediate conformation is a metastable state, as demonstrated by the short lifetime. Additionally, when starting from the intermediate conformation, the TVGG motif is more likely to relax to a random coil conformation.

Our results provide the following mechanistic picture. Folding of a peptide motif leads to a decrease in conformational entropy, which is compensated by favorable enthalpic interactions. For the folding kinetics of a small chain, such as the TVGG motif, conformational entropy is the likely factor that determines the energetic landscape ^39^. The barrier to partially folded *→* helix transformation is barrierless, yet in our simulations, the partially folded intermediate preferentially converts to the random coil conformation. This can be rationalized by considering the gain of conformational entropy for the partially folded *→* coil transformation. Additionally, in a folding pathway, the formation of intermediates can accelerate the kinetics ^40^.

We therefore propose that IP_6_ facilitates pentamer formation by accelerating the kinetics of the coil-to-helix transition mediated by an intermediate state during CA self-assembly.

## Discussion and Conclusions

The typical morphology of the canonical mature capsid is a fullerene cone. In accordance with Euler’s theorem, the HIV-1 capsid cone requires exactly 12 pentamers to ensure the closure of the polyhedron ^41^. The way the CA protein adjusts its structure to pentamers and hexamers for different regions of the lattice curvature remains an important area of investigation in HIV-1 structural biology. Analysis of the CA monomers from low-resolution structures of cores proposed that the interdomain hinge at the helix 9 likely provides the requisite flexibility to CA monomers to adapt to lattice curvature ^22,42^. High-resolution structures revealed that CA_NTD_ in hexamer and pentamer are nearly identical, with similar positioning of the hinge ^12^. Recent structural studies have identified the TVGG motif as a molecular switch that regulates hexamer or pentamer formation ^12,26^. CA tubes composed exclusively of hexamers are formed in abundance, while pentamers are rarely formed during spontaneous CA self-assembly ^18,43^. Therefore, CA monomers predominantly self-assemble to hexamers in the absence of IP_6_. In contrast, IP_6_ promotes the formation of cone-like structures^23^, which require pentamers for closure. Hence, a minimal combination of CA protein and IP_6_ is necessary for pentamer formation. Since structurally, the CA hexamers and pentamers in the mature capsid cone differ at the TVGG motif, IP_6_ likely triggers the TVGG conformational change.

After the formation of the pentamer during CA self-assembly^23^, the following molecular pathways are conceivable for the pentamer/hexamer switch: (1) coil-to-helix transition followed by IP_6_ binding at the pore locking the TVGG motif in the 3_10_ helix state, or (2) IP_6_ binding at the pore triggering the coil-to-helix transition. In this study, we tested both these pathways by simulating capsid patch with and without IP_6_. In the absence of IP_6_, the TVGG motif primarily adopted the coil conformation with no coil-to-helix transition. This likely explains the rarity of pentamers during self-assembly and eliminates the first pathway. In the presence of IP_6_, the likelihood of coil-to-helix transition increases, thus implicating IP_6_ as the primary trigger for the pentamer/hexamer switch. Our group’s previous atomistic simulation suggested that IP_6_ binding below the R18 ring is more favored in the pentamer than hexamer ^25^. The ligand density calculations of these simulation trajectories identified higher IP_6_ density below the R18 ring in the pentamer than in the hexamer. In our more extensive simulations of this paper, IP_6_ coordinated positively charged moieties from both R18 and K25 rings in the pentamer, leaving the coordination site above the R18 ring available for additional IP_6_ binding. This result is consistent with recent cryo-EM analysis that revealed a distinctly strong IP_6_ density below the R18 ring in the pentamer but a relatively weaker IP_6_ density in the hexamer ^24^. K25A mutants can form tubes, but their ability to assemble conical capsid is significantly diminished ^43^. If electrostatic repulsion within the central pore were the predominant factor in mature CA assembly, then K25A should exhibit enhanced conical assembly. In contrast, both the K25N mutant and wild-type CA can produce mature capsid in the presence of IP_6_^44^, suggesting a further role of IP_6_ binding at the K25 ring. In our simulations, the NTD-NTD interactions between helix 1-helix 2 are mediated by K25-M39 packing of adjacent subunits at the hydrophobic core. These molecular contacts play an important role in modulating the coil-to-helix transition consistent with the recently proposed ratchet mechanism of the hexamer/pentamer switch ^26^.

Our results so far have established the steps by which IP_6_ binding to the central pore triggers coil-to-helix conformational switching of the TVGG motif. IP_6_ binding allows K25-M39 packing at the hydrophobic core regulating coil-to-helix transition of the TVGG motif. The allosteric activation of protein due to ligand binding from the inactive to the active state is coupled with the change in the energetic landscape of the conformational change. Allosteric activation via ligand binding can occur through enthalpic regulation by modulation of the relative stability of the initial and final states^45^. In contrast, allosteric activation can occur through entropic regulation, where the relative stability of the end states remains unaltered^46^. Instead, additional intermediate conformations appear during the conformational change. Our results demonstrate that in the presence of IP_6_, an additional intermediate conformation regulates the coil-to-helix transition. Simultaneously, the barrier remains unchanged in the presence of IP_6_. Our results show that allostery between IP_6_ and TVGG motif conformational switching is entropically regulated.

To conclude, IP_6_ is necessary for the fullerene mode of assembly, while its absence results in the remodeling of pentameric defects into hexamer. The precise molecular mechanism underlying the insertion of a CA protomer into a transient metastable pentamer and the formation of a stable hexamer is of therapeutic interest. Our analysis reveals a high degree of fluctuation within the pentamer pore due to electropositive repulsion in the absence of IP_6_. Specifically, inter-protomer contact is weakened in helix 1 (movie S1), leading to the distortion of the compact pentameric molecular iris. A plausible mode of pentamer to hexamer remodeling during CA self-assembly can then be via the placement of a CA monomer at this defect site. Future investigations will require new, more highly resolved coarse-grained (CG) MD simulations, which can access timescales that are significantly longer than atomistic MD simulations in this work and can better elucidate the mechanistic pathways of pentamer to hexamer remodeling.

## Methods and Materials

### System and simulation setup

The initial fully atomistic structures of the CA hexamer and pentamers (residues 1-220) were constructed from the cryo-electron tomography structures PDB: 5MCX and PDB: 5MCY, respectively^47^. These atomic structures are derived from intact virus particles. Residues 220-231 were obtained from NMR structure of CA_CTD_ (PDB: 2KOD) ^48^. The atomistic structures of CA hexamer and pentamers were aligned to an equilibrated atomistic structure of a capsid cone ^34^ to construct a capsid patch with a central pentamer and five surrounding hexamers. In the IP_6_ simulations, six IP_6_ were positioned roughly 3 Å above the R18 pore in each capsomer based on X-ray crystal structure ^18^. The protein-cofactor complex was placed in the center of the simulation cell such that the distance between the edge of the protein and the box boundary is 1.2 nm. The protein-cofactor complex was solvated by 795,958 water molecules andthe salt concentrations were adjusted to maintain 150 mM NaCl and produce a charge-neutral system, resulting in a total of 2,600,199 atoms.

Energy minimization was performed using steepest descent algorithm, and the minimization was terminated when the maximum force was smaller than 1000.0 kJ/mol/nm. The composite system was then equilibrated in constant NVT ensemble by applying harmonic positional restraint (spring constant value of 1000 kJ/mol/nm^2^) on protein heavy atoms for 10 ns. The temperature of the system was maintained at 310 K using stochastic velocity rescaling thermostat with a time constant of 1 ps ^49^. The harmonic positional restraints were removed, and the system was equilibrated in constant NPT ensemble at 310 K and 1 bar for an additional 100 ns. The configurations at the endpoint of the NPT equilibration were used as initial configurations for the production runs. The production runs were performed at 310 K and 1 bar. In the NPT simulations, the temperature was maintained using Nose-Hoover chain thermostat with a 2 ps time constant and pressure with Parrinello-Rahman barostat with a 10 ps time constant ^50,51^.

We prepared a partial capsid patch containing a central CA pentamer and 2 closest CA monomers of each hexamer surrounding the pentamer. The partial capsid patch was used to perform conformational sampling simulations and WT-MetaD simulations. In these simulations, the CA monomers that are part of the hexameric lattice by applying harmonic positional restraint (spring constant value of 1000 kJ/mol/nm^2^). The solvated partial capsid patch system was prepared with the identical protocol as the full capsid patch system. The system was solvated by 274,274 water molecules. Salt concentrations were adjusted to maintain 150 mM NaCl, resulting in a system with a total of 878,428 atoms.

All MD simulations were performed with periodic boundary conditions in all directions. The protein was modeled with CHARMM36m force field ^52^, and water was modeled with TIP3P parameters ^53^. The parameters of IP_6_ were generated with cgenff ^54^. The bonds between the protein heavy atoms and hydrogen atoms were constrained with LINCS algorithm. Electrostatic interactions were computed using the particle mesh Ewald method with a cutoff of 1.2 nm ^55^. The van der Waals force was truncated smoothly to zero between 1.0 and 1.2 nm. The equations of motion were integrated using the velocity Verlet algorithm with a timestep of 2 fs ^56^. The unbiased simulations were performed using GROMACS version 2019.6 ^57^. All restrained simulations and WT-MetaD simulations were performed using GROMACS version 2020.4 patched with PLUMED 2.6 ^58^.

### Well-tempered metadynamics simulation setup

We estimated the free energy landscape of the coil-to-helix transition of the TVGG motif with and without IP_6_ bound at the central pore using WT-MetaD simulations ^38^. We used two collective variables (CVs) to perform our WT-MetaD simulation: (1) the distances between the center of mass (COM) of the side chain heavy atoms of K25 of *CA*_1_ and M39 of *CA*_2_ denoted as *d*_*K25-M39*_, and (2) helicity parameter (*h*_*V*59_) of V59 residue of *CA*_1_. Here, *CA*_1_ and *CA*_2_ are two adjacent monomers in a pentamer. The helicity parameter (*h*_*V*59_) is defined as 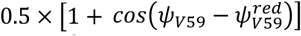. Here, ψ_*V*59_ is the dihedral angle centered at the V59 residue of *CA*_1_ 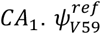 is the reference dihedral angle that corresponds to the value of the ideal α-helix. The selection of the CVs was informed by unbiased atomistic simulation results.

We estimated the free energy profile of coil-to-helix transition using the approach in ref. ^33^. Briefly, the sampled space in WT-MetaD with microscopic coordinate *R* is associated with a reweighting factor *w*(*R*_1_ = *e*^*V(s(R),t)-c(t)*^. Here *V*(*x)* is the bias *c*(*t)* is referred to as a time-dependent bias offset. The term *c*(*t)* is estimated “on the fly” in the MetaD simulations. In our simulations, the biasing factor was set to 10, the bias height was set to 0.2 kcal/mol, and the biasing frequency was set to 5000 timesteps. These parameters were chosen to ensure that the biasing potential was not added too aggressively. The variances of Gaussian bias were set to be 0.05 Å and 0.05 for *d*_*K25-M39*_ and *h*_*V*59_, respectively. The upper bound of the *d*_*K25-M39*_ was set to 14 Å.

### Analysis of MD simulations

In the production runs, the configuration of the system was saved every 500 ps interval. MDAnalysis package^59^ was used to access and align the trajectories. Positions of the C_α_ center of mass of the relevant residues were used for the calculation of inter-subunit distances. The average distance of all (i, i+1) pairs within the pentamer was used for the contact maps. Density of IP_6_ was calculated by mapping the atomic densities of the ligand on a discretized grid with a volume of 0.5 Å × 0.5 Å × 0.5 Å.

## Supporting information

Supporting Information for Kinetic Implications of IP6 Anion Binding on the Molecular Switch of the HIV-1 Capsid Assembly

## Acknowledgements

This research was supported in part by the National Institute of Allergy and Infectious Diseases (NIAID) of the National Institutes of Health (NIH) grant R01AI178550 to GAV. The content is solely the responsibility of the authors and does not necessarily represent the official views of the National Institutes of Health. Partial support was also received from the Frost Institute for Chemistry and Molecular Science at the University of Miami to MY. Simulations were performed using resources by the Advanced Cyberinfrastructure Coordination Ecosystem: Services & Support program, which is supported by NSF grants numbers 2138259, 2138286, 2138307, 2137603, and 2138296, and Frontera ^60^ (at TACC) funded by the NSF (OAC-1818253). We thank Dr. Yihang Wang for helpful discussions.

## References

1. E. O. Freed. “HIV-1 assembly, release and maturation”, Nat. Rev. Microbiol. 13, 484–496 (2015).

2. W. I. Sundquist and H. G. Krausslich. “HIV-1 assembly, budding, and maturation”, Cold Spring Harb. Perspect. Med. 2, a006924 (2012).

3. B. K. Ganser-Pornillos, M. Yeager and O. Pornillos. “Assembly and architecture of HIV”, Adv. Exp. Med. Biol. 726, 441–465 (2012).

4. J. A. Briggs and H.G. Kräusslich. “The molecular architecture of HIV”, J. Mol. Biol. 410, 491–500 (2011).

5. J. Rasaiyaah, C. P. Tan, A. J. Fletcher, A. J. Price, C. Blondeau, L. Hilditch, D. A. Jacques, D. L. Selwood, L. C. James, M. Noursadeghi and G. J. Towers. “HIV-1 evades innate immune recognition through specific cofactor recruitment”, Nature 503, 402–405 (2013).

6. V. Zila, E. Margiotta, B. Turoňová, T. G. Müller, C. E. Zimmerli, S. Mattei, M. Allegretti, K. Börner, J. Rada and B. Müller. “Cone-shaped HIV-1 capsids are transported through intact nuclear pores”, Cell 184, 1032-1046. e1018 (2021).

7. A. Dharan, N. Bachmann, S. Talley, V. Zwikelmaier and E. M. Campbell. “Nuclear pore blockade reveals that HIV-1 completes reverse transcription and uncoating in the nucleus”, Nature microbiology 5, 1088–1095 (2020).

8. A. Hudait and G. A. Voth. “HIV-1 capsid shape, orientation, and entropic elasticity regulate translocation into the nuclear pore complex”, Proceedings of the National Academy of Sciences 121, e2313737121 (2024).

9. T. G. Müller, V. Zila, K. Peters, S. Schifferdecker, M. Stanic, B. Lucic, V. Laketa, M. Lusic, B. Müller and H.-G. Kräusslich. “HIV-1 uncoating by release of viral cDNA from capsid-like structures in the nucleus of infected cells”, eLife 10, e64776 (2021).

10. R. C. Burdick, C. Li, M. Munshi, J. M. O. Rawson, K. Nagashima, W.-S. Hu and V. K. Pathak. “HIV-1 uncoats in the nucleus near sites of integration”, Proceedings of the National Academy of Sciences 117, 5486–5493 (2020).

11. C. Li, R. C. Burdick, K. Nagashima, W.-S. Hu and V. K. Pathak. “HIV-1 cores retain their integrity until minutes before uncoating in the nucleus”, Proceedings of the National Academy of Sciences 118, e2019467118 (2021).

12. J. C. V. Stacey, A. Tan, J. M. Lu, L. C. James, R. A. Dick and J. A. G. Briggs. “Two structural switches in HIV-1 capsid regulate capsid curvature and host factor binding”, Proceedings of the National Academy of Sciences 120, e2220557120 (2023).

13. T. Ni, S. Gerard, G. Zhao, K. Dent, J. Ning, J. Zhou, J. Shi, J. Anderson-Daniels, W. Li, S. Jang, A. N. Engelman, C. Aiken and P. Zhang. “Intrinsic curvature of the HIV-1 CA hexamer underlies capsid topology and interaction with cyclophilin A”, Nature Structural & Molecular Biology 27, 855–862 (2020).

14. L. Deshmukh, C. D. Schwieters, A. Grishaev, R. Ghirlando, J. L. Baber and G. M. Clore. “Structure and Dynamics of Full-Length HIV-1 Capsid Protein in Solution”, J.Am.Chem.Soc. 135, 16133–16147 (2013).

15. A. T. Gres, K. A. Kirby, V. N. KewalRamani, J. J. Tanner, O. Pornillos and S. G. Sarafianos. “X-ray crystal structures of native HIV-1 capsid protein reveal conformational variability”, Science 349, 99–103 (2015).

16. O. Pornillos, B. K. Ganser-Pornillos and M. Yeager. “Atomic-level modelling of the HIV capsid”, Nature 469, 424-427 %428 Jan 420 %! Atomic-level modelling of the HIV capsid %@ 1476-4687 (2011).

17. R. A. Dick, D. L. Mallery, V. M. Vogt and L. C. James. “IP6 Regulation of HIV Capsid Assembly, Stability, and Uncoating”, Viruses 10 (2018).

18. R. A. Dick, K. K. Zadrozny, C. Xu, F. K. M. Schur, T. D. Lyddon, C. L. Ricana, J. M. Wagner, J. R. Perilla, B. K. Ganser-Pornillos, M. C. Johnson, O. Pornillos and V. M. Vogt. “Inositol phosphates are assembly co-factors for HIV-1”, Nature 560, 509–512 (2018).

19. D. L. Mallery, K. M. R. Faysal, A. Kleinpeter, M. S. C. Wilson, M. Vaysburd, A. J. Fletcher, M. Novikova, T. Böcking, E. O. Freed, A. Saiardi and L. C. James. “Cellular IP6 Levels Limit HIV Production while Viruses that Cannot Efficiently Package IP6 Are Attenuated for Infection and Replication”, Cell Reports 29, 3983-3996.e3984 (2019).

20. S. Campbell, R. J. Fisher, E. M. Towler, S. Fox, H. J. Issaq, T. Wolfe, L. R. Phillips and A. Rein. “Modulation of HIV-like particle assembly in vitro by inositol phosphates”, Proceedings of the National Academy of Sciences 98, 10875–10879 (2001).

21. D. A. Jacques, W. A. McEwan, L. Hilditch, A. J. Price, G. J. Towers and L. C. James. “HIV-1 uses dynamic capsid pores to import nucleotides and fuel encapsidated DNA synthesis”, Nature 536, 349–353 (2016).

22. T. Ni, Y. Zhu, Z. Yang, C. Xu, Y. Chaban, T. Nesterova, J. Ning, T. Böcking, M. W. Parker, C. Monnie, J. Ahn, J. R. Perilla and P. Zhang. “Structure of native HIV-1 cores and their interactions with IP6 and CypA”, Science Advances 7, eabj5715 (2021).

23. M. Gupta, A. J. Pak and G. A. Voth. “Critical mechanistic features of HIV-1 viral capsid assembly”, Science Advances 9, eadd7434 (2023).

24. C. M. Highland, A. Tan, C. L. Ricaña, J. A. G. Briggs and R. A. Dick. “Structural insights into HIV-1 polyanion-dependent capsid lattice formation revealed by single particle cryo-EM”, Proceedings of the National Academy of Sciences 120, e2220545120 (2023).

25. A. Yu, E. M. Y. Lee, J. Jin and G. A. Voth. “Atomic-scale characterization of mature HIV-1 capsid stabilization by inositol hexakisphosphate (IP6)”, Sci. Adv. 6 (2020).

26. R. T. Schirra, N. F. B. Dos Santos, K. K. Zadrozny, I. Kucharska, B. K. Ganser-Pornillos and O. Pornillos. “A molecular switch modulates assembly and host factor binding of the HIV-1 capsid”, Nature structural & molecular biology 30, 383–390 (2023).

27. O. Pornillos, B. K. Ganser-Pornillos, B. N. Kelly, Y. Hua, F. G. Whitby, C. D. Stout, W. I. Sundquist, C. P. Hill and M. Yeager. “X-ray structures of the hexameric building block of the HIV capsid”, Cell 137, 1282–1292 (2009).

28. M. G. Mateu. “The capsid protein of human immunodeficiency virus: intersubunit interactions during virus assembly”, The FEBS Journal 276, 6098–6109 (2009).

29. J.-P. Changeux. “Allostery and the Monod-Wyman-Changeux Model After 50 Years”, Annual Review of Biophysics 41, 103–133 (2012).

30. A. Cooper and D. T. F. Dryden. “Allostery without conformational change”, European Biophysics Journal 11, 103–109 (1984).

31. A. Barducci, G. Bussi and M. Parrinello. “Well-Tempered Metadynamics: A Smoothly Converging and Tunable Free-Energy Method”, Physical Review Letters 100, 020603 (2008).

32. J. F. Dama, M. Parrinello and G. A. Voth. “Well-Tempered Metadynamics Converges Asymptotically”, Physical Review Letters 112, 240602 (2014).

33. P. Tiwary and M. Parrinello. “A Time-Independent Free Energy Estimator for Metadynamics”, The Journal of Physical Chemistry B 119, 736–742 (2015).

34. A. Yu, E. M. Y. Lee, J. A. G. Briggs, B. K. Ganser-Pornillos, O. Pornillos and G. A. Voth. “Strain and rupture of HIV-1 capsids during uncoating”, Proceedings of the National Academy of Sciences 119, e2117781119 (2022).

35. A. T. Gres, K. A. Kirby, W. M. McFadden, H. Du, D. Liu, C. Xu, A. J. Bryer, J. R. Perilla, J. Shi, C. Aiken, X. Fu, P. Zhang, A. C. Francis, G. B. Melikyan and S. G. Sarafianos. “Multidisciplinary studies with mutated HIV-1 capsid proteins reveal structural mechanisms of lattice stabilization”, Nature Communications 14, 5614 (2023).

36. A. Bhattacharya, S. L. Alam, T. Fricke, K. Zadrozny, J. Sedzicki, A. B. Taylor, B. Demeler, O. Pornillos, B. K. Ganser-Pornillos and F. Diaz-Griffero. “Structural basis of HIV-1 capsid recognition by PF74 and CPSF6”, Proceedings of the National Academy of Sciences 111, 18625–18630 (2014).

37. L. Lamorte, S. Titolo, C. T. Lemke, N. Goudreau, J.-F. Mercier, E. Wardrop, V. B. Shah, U. K. von Schwedler, C. Langelier and S. S. R. Banik. “Discovery of novel small-molecule HIV-1 replication inhibitors that stabilize capsid complexes”, Antimicrobial agents and chemotherapy 57, 4622–4631 (2013).

38. O. Valsson, P. Tiwary and M. Parrinello. “Enhancing Important Fluctuations: Rare Events and Metadynamics from a Conceptual Viewpoint”, (2016).

39. A. R. Dinner, A. Šali, L. J. Smith, C. M. Dobson and M. Karplus. “Understanding protein folding via free-energy surfaces from theory and experiment”, Trends in Biochemical Sciences 25, 331–339 (2000).

40. C. Wagner and T. Kiefhaber. “Intermediates can accelerate protein folding”, Proceedings of the National Academy of Sciences 96, 6716–6721 (1999).

41. S. Li, C. P. Hill, W. I. Sundquist and J. T. Finch. “Image reconstructions of helical assemblies of the HIV-1 CA protein”, Nature 407, 409–413 (2000).

42. G. Zhao, J. R. Perilla, E. L. Yufenyuy, X. Meng, B. Chen, J. Ning, J. Ahn, A. M. Gronenborn, K. Schulten, C. Aiken and P. Zhang. “Mature HIV-1 capsid structure by cryo-electron microscopy and all-atom molecular dynamics”, Nature 497, 643–646 (2013).

43. N. Renner, D. L. Mallery, K. M. R. Faysal, W. Peng, D. A. Jacques, T. Böcking and L. C. James. “A lysine ring in HIV capsid pores coordinates IP6 to drive mature capsid assembly”, PLOS Pathogens 17, e1009164 (2021).

44. C. Xu, D. K. Fischer, S. Rankovic, W. Li, R. A. Dick, B. Runge, R. Zadorozhnyi, J. Ahn, C. Aiken, T. Polenova, A. N. Engelman, Z. Ambrose, I. Rousso and J. R. Perilla. “Permeability of the HIV-1 capsid to metabolites modulates viral DNA synthesis”, PLOS Biology 18, e3001015 (2020).

45. M. F. Perutz. “Stereochemistry of Cooperative Effects in Haemoglobin: Haem–Haem Interaction and the Problem of Allostery”, Nature 228, 726–734 (1970).

46. S. E. Reichheld, Z. Yu and A. R. Davidson. “The induction of folding cooperativity by ligand binding drives the allosteric response of tetracycline repressor”, Proceedings of the National Academy of Sciences 106, 22263–22268 (2009).

47. S. Mattei, B. Glass, W. J. Hagen, H.G. Kräusslich and J. A. Briggs. “The structure and flexibility of conical HIV-1 capsids determined within intact virions”, Science 354, 1434–1437 (2016).

48. I. J. Byeon, X. Meng, J. Jung, G. Zhao, R. Yang, J. Ahn, J. Shi, J. Concel, C. Aiken, P. Zhang and A. M. Gronenborn. “Structural convergence between Cryo-EM and NMR reveals intersubunit interactions critical for HIV-1 capsid function”, Cell 139, 780–790 (2009).

49. G. Bussi, D. Donadio and M. Parrinello. “Canonical sampling through velocity rescaling”, The Journal of Chemical Physics 126, 014101 (2007).

50. D. J. Evans and B. L. Holian. “The Nose–Hoover thermostat”, The Journal of Chemical Physics 83, 4069–4074 (1985).

51. M. Parrinello and A. Rahman. “Polymorphic transitions in single crystals: A new molecular dynamics method”, Journal of Applied Physics 52, 7182–7190 (1981).

52. J. Huang, S. Rauscher, G. Nawrocki, T. Ran, M. Feig, B. L. de Groot, H. Grubmüller and A. D. MacKerell. “CHARMM36m: an improved force field for folded and intrinsically disordered proteins”, Nature Methods 14, 71–73 (2017).

53. W. L. Jorgensen, J. Chandrasekhar, J. D. Madura, R. W. Impey and M. L. Klein. “Comparison of simple potential functions for simulating liquid water”, The Journal of Chemical Physics 79, 926–935 (1983).

54. K. Vanommeslaeghe, E. Hatcher, C. Acharya, S. Kundu, S. Zhong, J. Shim, E. Darian, O. Guvench, P. Lopes, I. Vorobyov and A. D. Mackerell Jr. “CHARMM general force field: A force field for drug-like molecules compatible with the CHARMM all-atom additive biological force fields”, Journal of Computational Chemistry 31, 671–690 (2010).

55. T. Darden, D. York and L. Pedersen. “Particle mesh Ewald: An N·log(N) method for Ewald sums in large systems”, The Journal of Chemical Physics 98, 10089–10092 (1993).

56. W. C. Swope, H. C. Andersen, P. H. Berens and K. R. Wilson. “A computer simulation method for the calculation of equilibrium constants for the formation of physical clusters of molecules: Application to small water clusters”, The Journal of Chemical Physics 76, 637–649 (1982).

57. M. J. Abraham, T. Murtola, R. Schulz, S. Páll, J. C. Smith, B. Hess and E. Lindahl. “GROMACS: High performance molecular simulations through multi-level parallelism from laptops to supercomputers”, SoftwareX 1-2, 19–25 (2015).

58. G. A. Tribello, M. Bonomi, D. Branduardi, C. Camilloni and G. Bussi. “PLUMED 2: New feathers for an old bird”, Computer Physics Communications 185, 604–613 (2014).

59. N. Michaud-Agrawal, E. J. Denning, T. B. Woolf and O. Beckstein. “MDAnalysis: a toolkit for the analysis of molecular dynamics simulations”, Journal of computational chemistry 32, 2319–2327 (2011).

60. D. Stanzione, J. West, R. T. Evans, T. Minyard, O. Ghattas and D. K. Panda. in Practice and Experience in Advanced Research Computing 106–111 (2020).

