## Supporting Information for Kinetic Implications of IP6 Anion Binding on the Molecular Switch of the HIV-1 Capsid Assembly for "Kinetic Implications of IP_6_ Anion Binding on the Molecular Switch of the HIV-1 Capsid Assembly"

[#] equal contribution

The PDF file contains:

Figs. S1 to S16
Legends for movies S1 and S2

Here, we present results from additional unbiased all-atom simulations of the capsid patch under $-IP_6$ and $+ IP_6$ conditions. Three replicas were simulated for each case. Without $IP_6$, the TVGG motif primarily stayed in coil configuration, only transiently exploring the folded conformations. Notably, the central pore is more dynamic in the absence of $IP_6$, resulting in the relocation of M39 residues towards the 'pawl' state near the TVGG motif, as is evident from the K25-M39 and T58-M39 distance analyses. In the presence of $IP_6$, M39 preferred to stay in the 'ratchet' state packing against V24/K25, which led to occasional folding of the TVGG motif into a partial $3_{10}$-helix conformation.

**Replica 1 ($-IP_6$)**

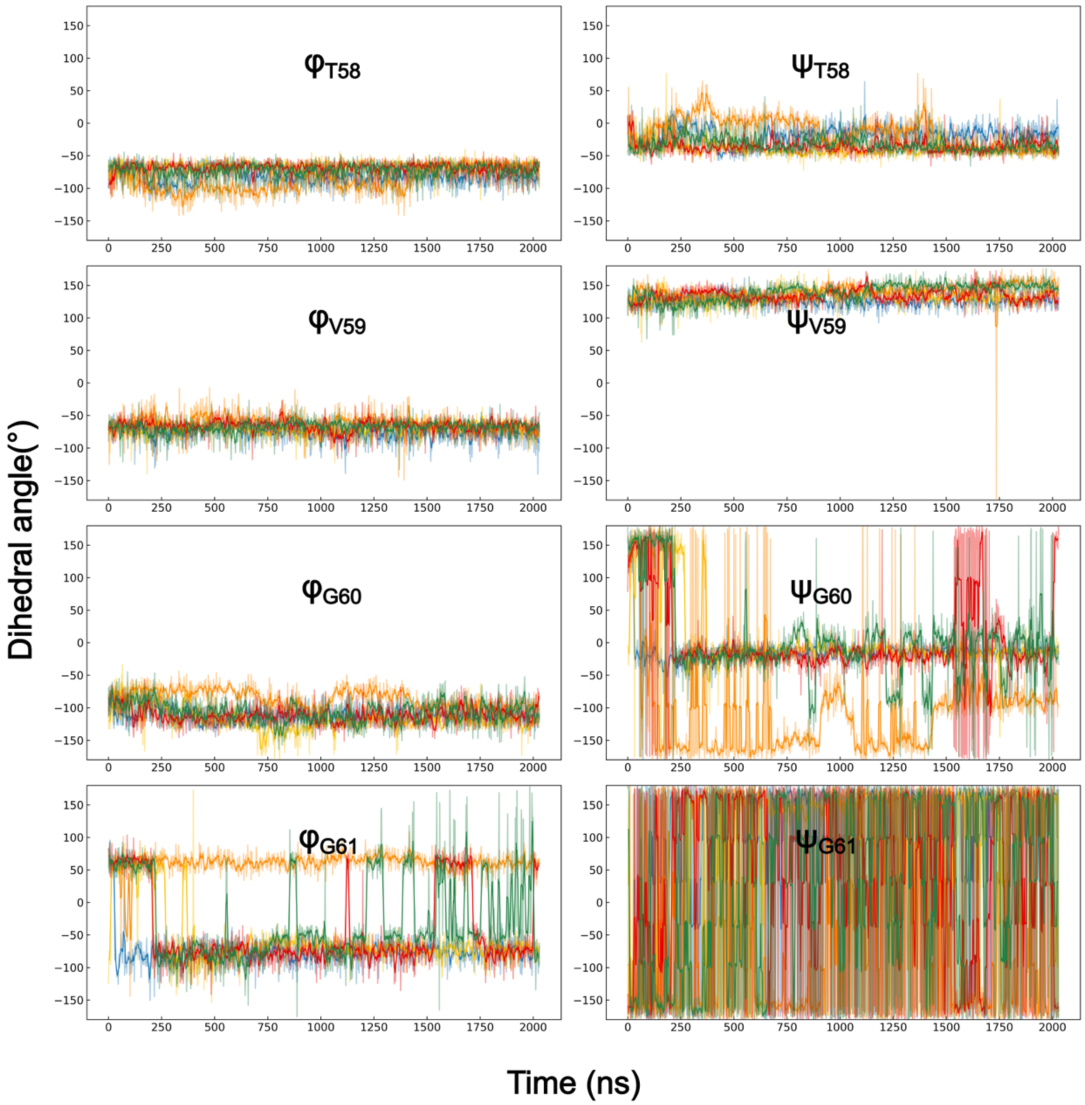


**Fig. S1.** Time series of the TVGG backbone dihedral angles in the absence of $IP_6$ ($-IP_6$). T58 residues in the capsid pentamer adopted a folded configuration except for the T58 residue in the γ-domain, which eventually converged to a $3_{10}$-helix conformation. V59, G60 residues were ordered but deviated from a typical folded conformation. G61 residues were relatively more mobile.

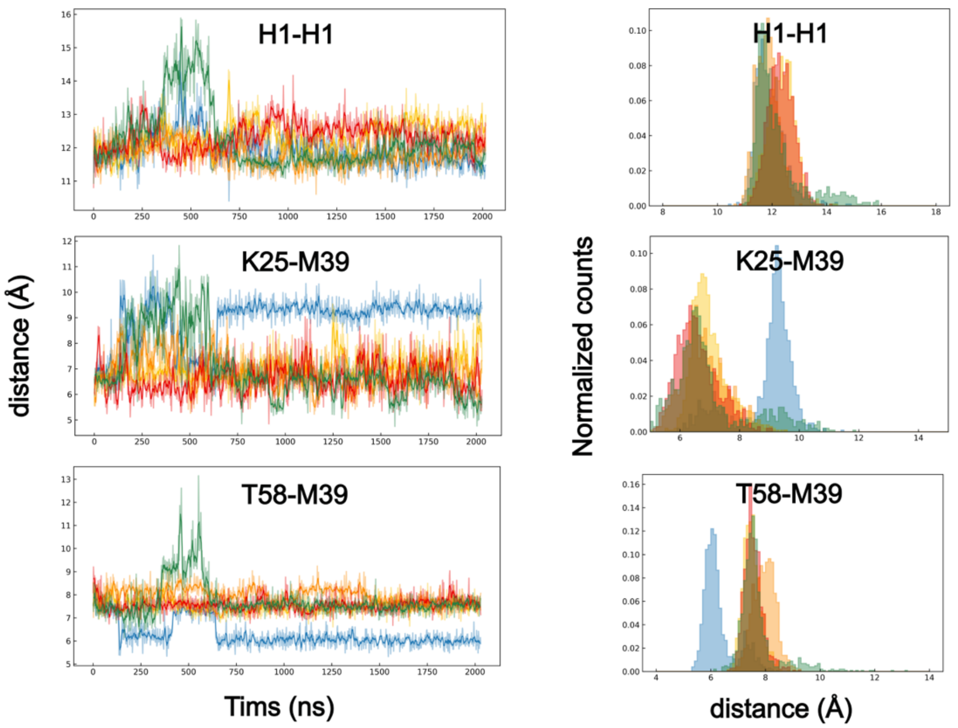


**Fig. S2.** Time series of the capsid structural network that links $IP_6$ to the TVGG motif in the absence of $IP_6$ (-$IP_6$). In this case, the central pore was relatively ordered. In the pentamer, a stable K25/M39 packing was observed between NTD of adjacent CA monomers. However, M39 residue in the β-domain moved away from α-domain V24/K25, breaking the 'ratchet' state preferred in pentamers.

**Replica 2 (-IP$_6$)**

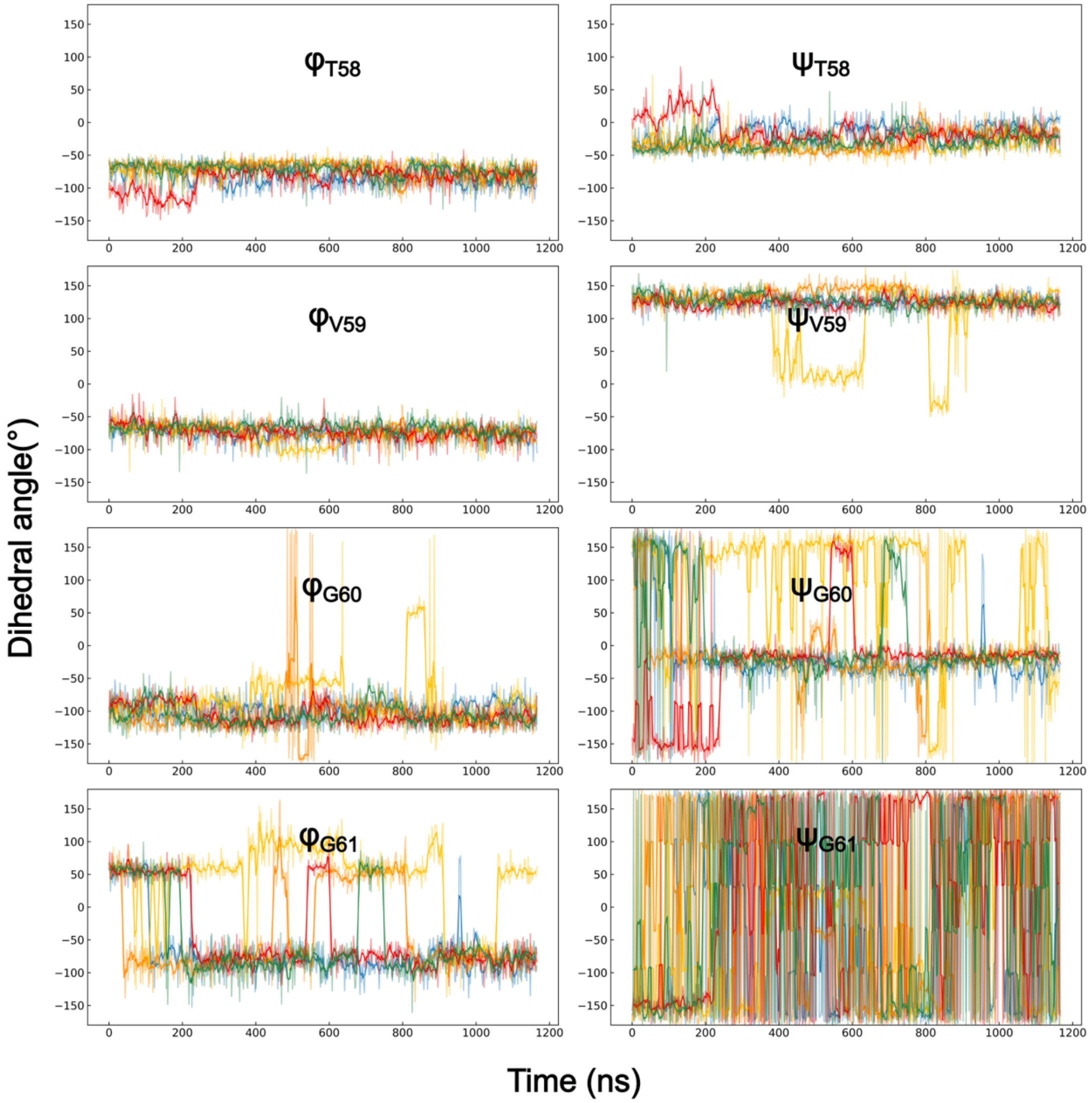


**Fig. S3.** Time series of the TVGG backbone dihedral angles in the absence of IP$_6$ (-IP$_6$). β-domain V59 residue explored folded conformations transiently, however, all chains converged to coil conformation.

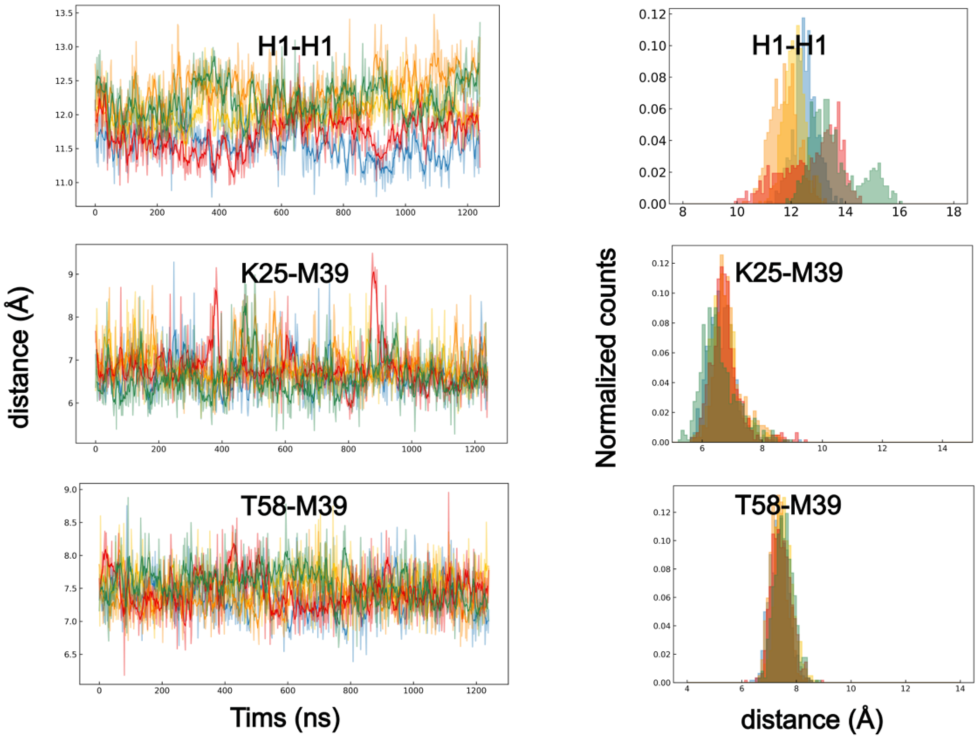


**Fig. S4.** Time series of the capsid structural network that links $IP_6$ to the TVGG motif in the absence of $IP_6$ (-$IP_6$). The capsid pentamer stayed in the 'ratchet' conformation during the time of the simulation which allowed the TVGG motif in the β-domain (Fig. S3) to explore helix-like folded conformations.

**Replica 3 (-IP$_6$)**

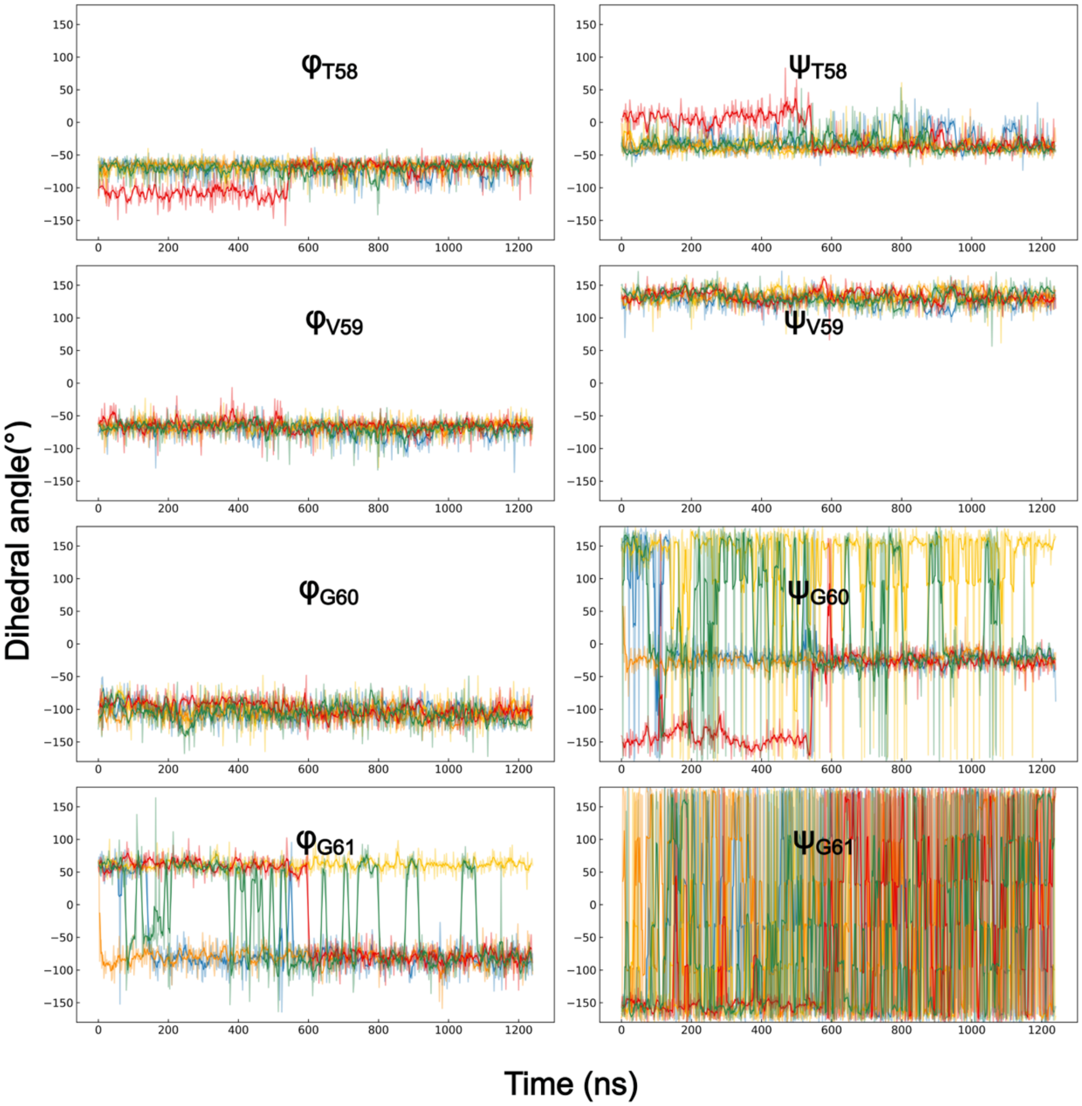


**Fig. S5.** Time series of the TVGG backbone dihedral angles in the absence of IP$_6$ (-IP$_6$).

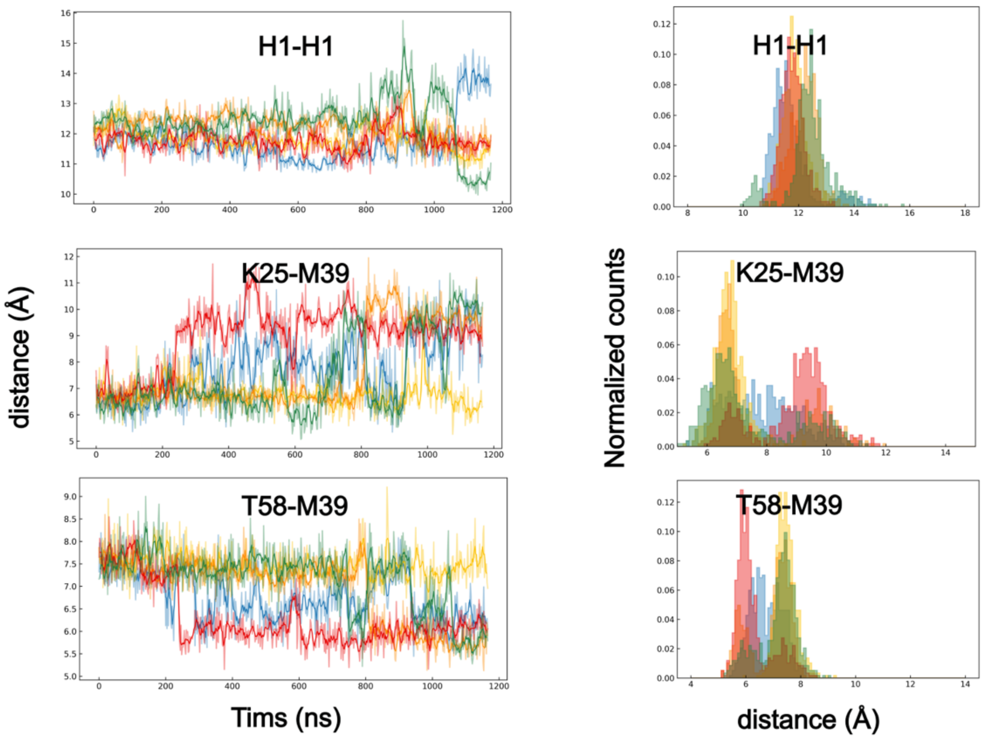


**Fig. S6.** Time series of the capsid structural network that links $IP_6$ to the TVGG motif in the absence of $IP_6$ ($-IP_6$). The central pentamer pore was disordered, leading to the deviation of M39 residues from the 'pawl' conformation in multiple CA domains.

**Replica 1 (+$IP_6$)**

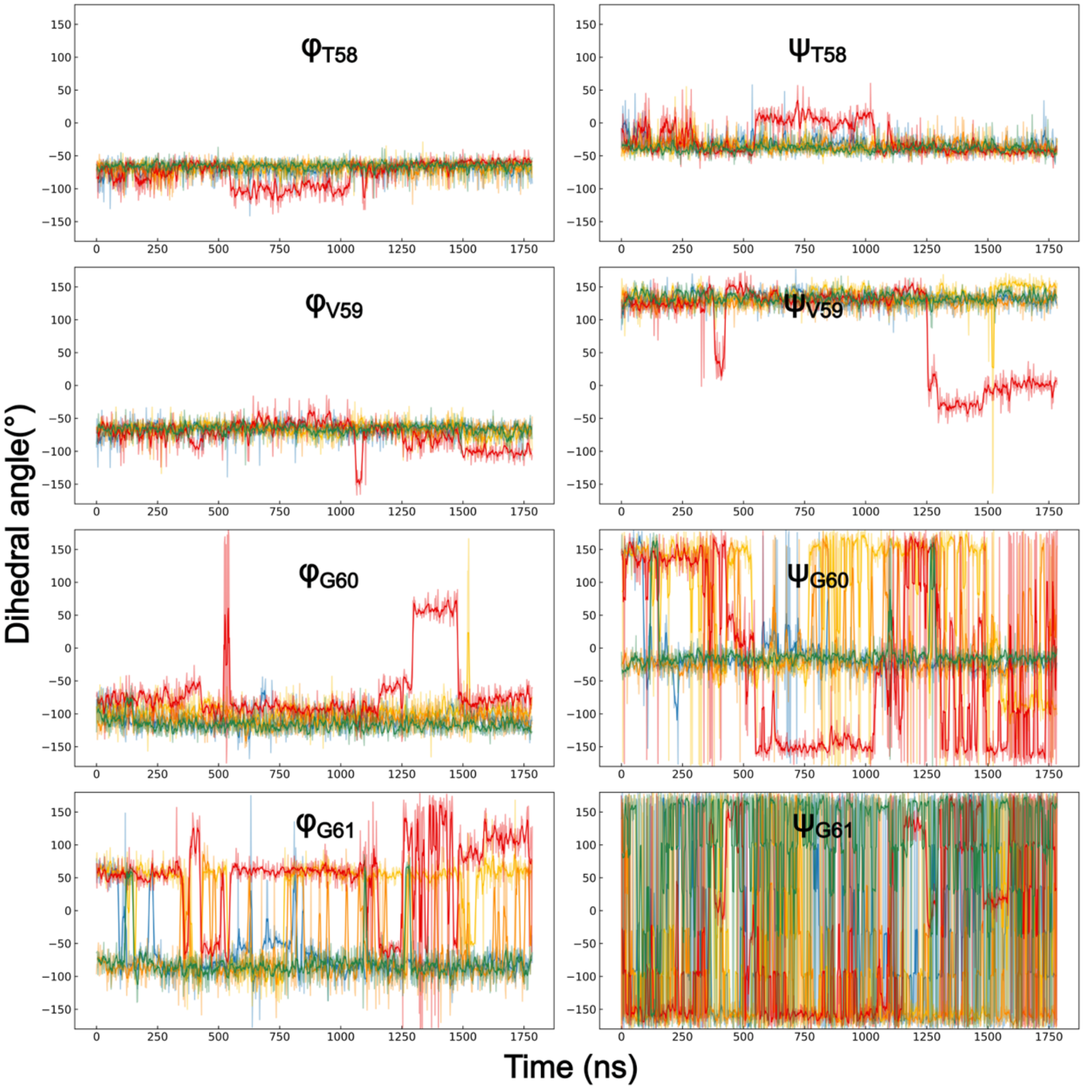


**Fig. S7.** Time series of the TVGG backbone dihedral angles in the presence of $IP_6$ (+$IP_6$). In this replica, T58 residue in the $\alpha$-domain CA unfolded briefly, following which T58 and V59 residues switched to a folded conformation, forming a partial $3_{10}$-helix structure.

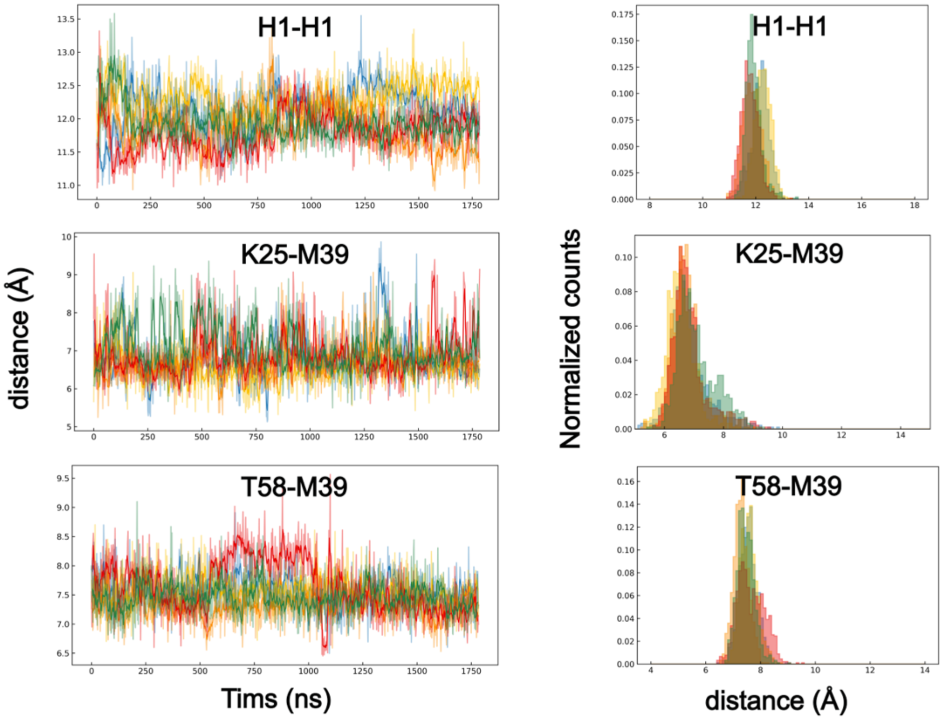


**Fig. S8.** Time series of the capsid structural network that links $IP_6$ to the TVGG motif in the presence of $IP_6$ (+$IP_6$). $IP_6$ stabilized the electrostatic repulsion within the pentamer central pore. This led to an ordered K25 ring accommodating stable packing with M39 of adjacent CA domains. The pentamer did not deviate from 'pawl' conformation during the simulation.

**Replica 2 (+$IP_6$)**

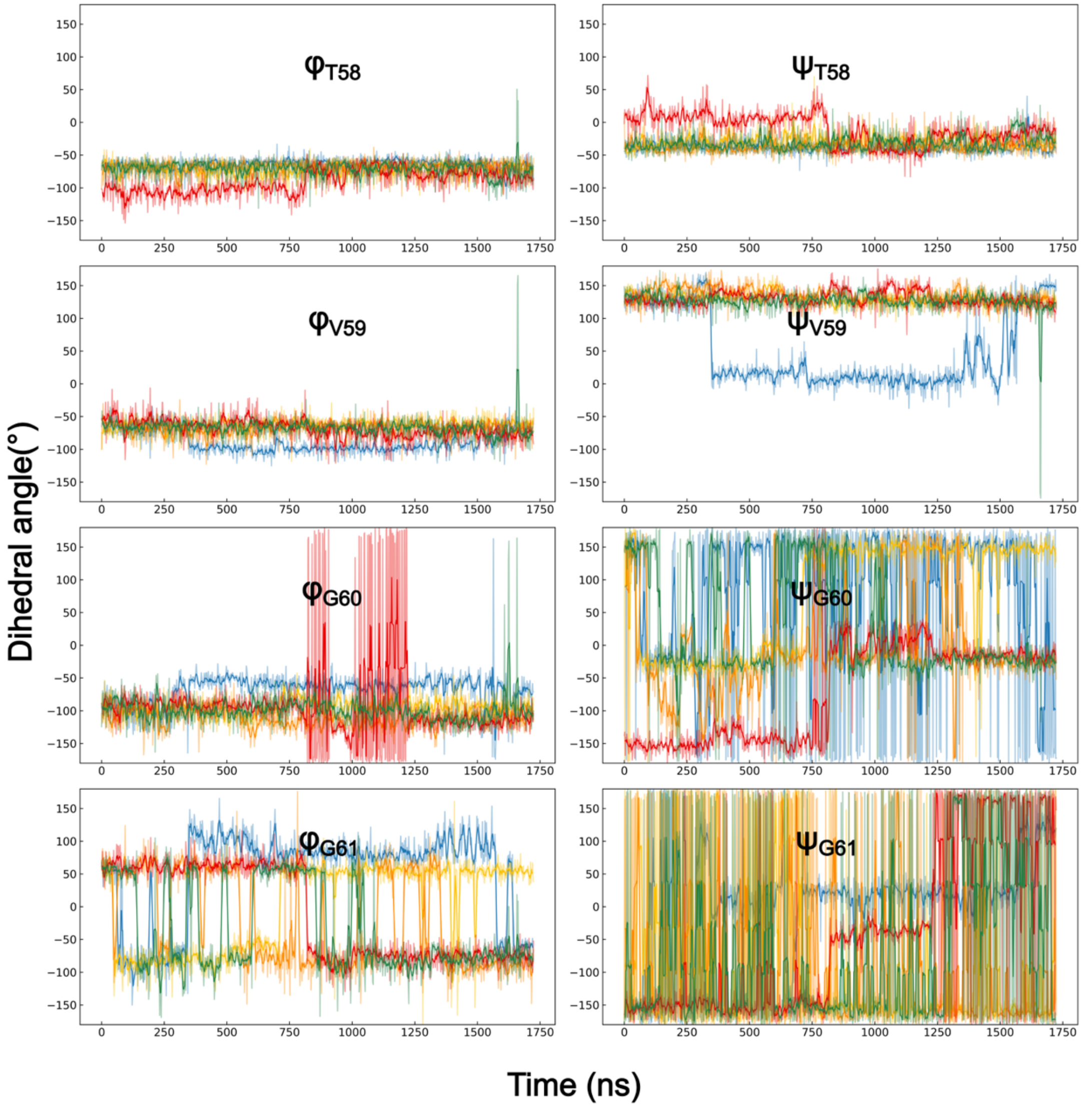


**Fig. S9.** Time series of the TVGG backbone dihedral angles in the presence of $IP_6$ (+$IP_6$).

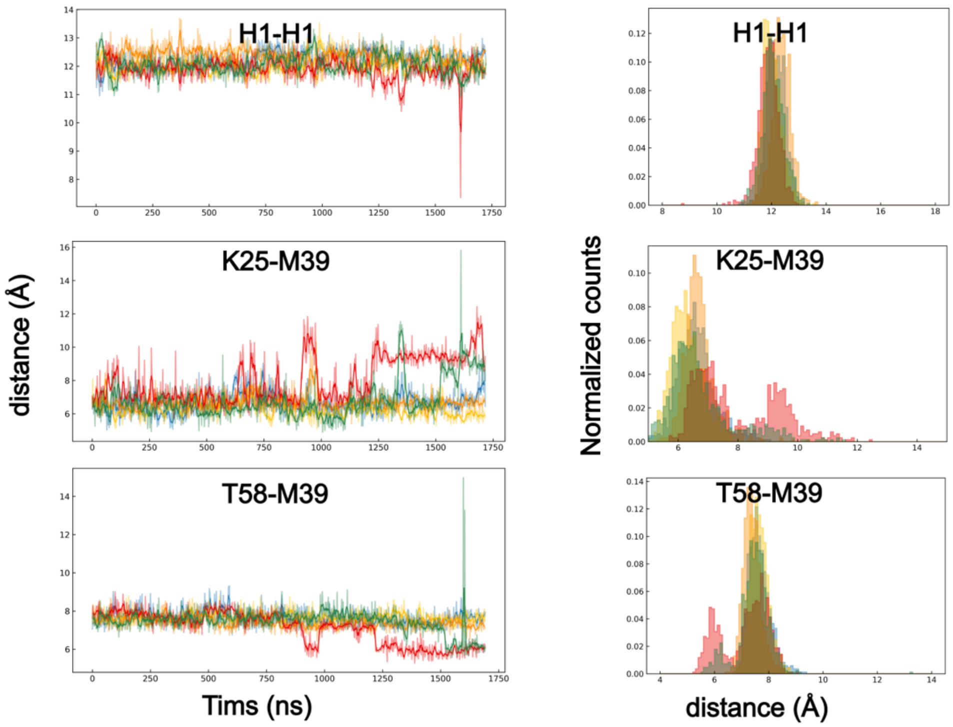


**Fig. S10.** Time series of the capsid structural network that links $IP_6$ to the TVGG motif in the presence of $IP_6$ (+$IP_6$).

**Replica 3 (+$IP_6$)**

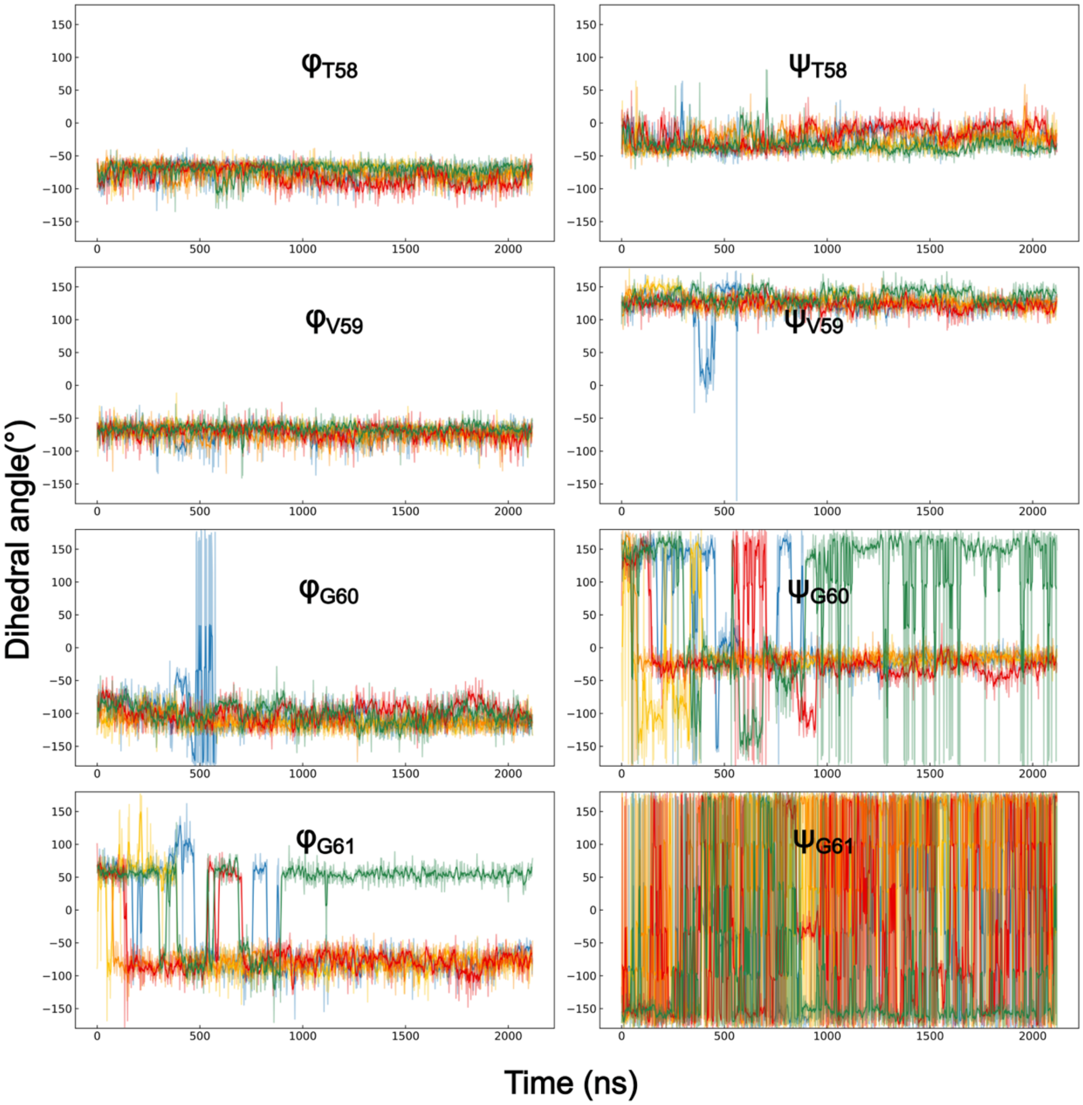


**Fig. S11.** Time series of the TVGG backbone dihedral angles in the presence of $IP_6$ (+$IP_6$).

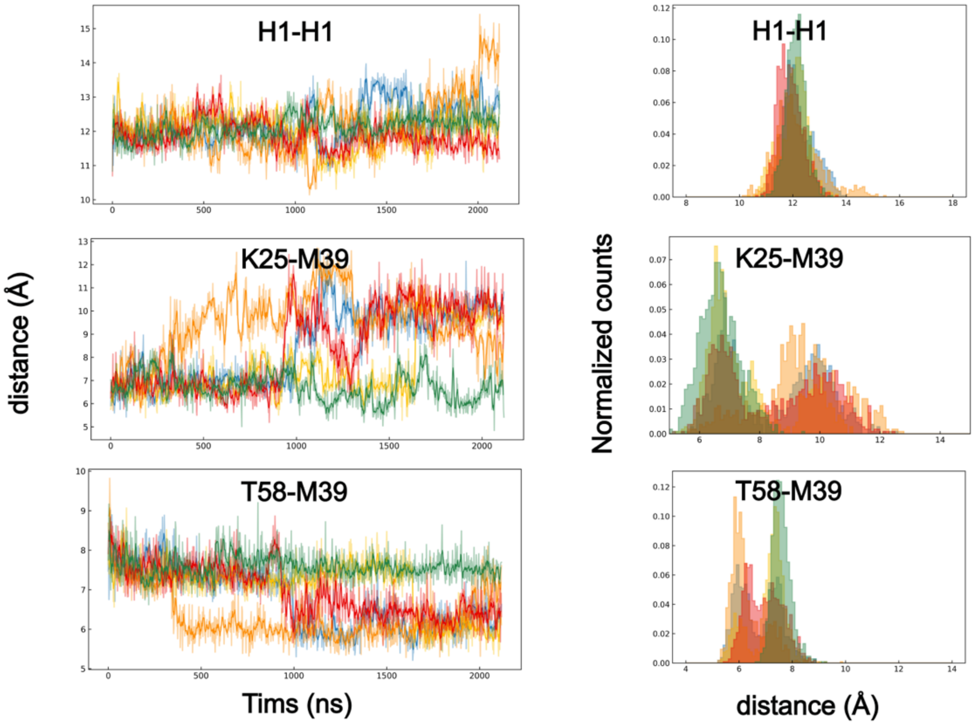


**Fig. S12.** Time series of the capsid structural network that links $IP_6$ to the TVGG motif in the presence of $IP_6$ (+$IP_6$). No major coil-to-helix transition was observed in this case (Fig. S11) since multiple CA domains in the $IP_6$-bound pentamer slipped into the 'ratchet' state.

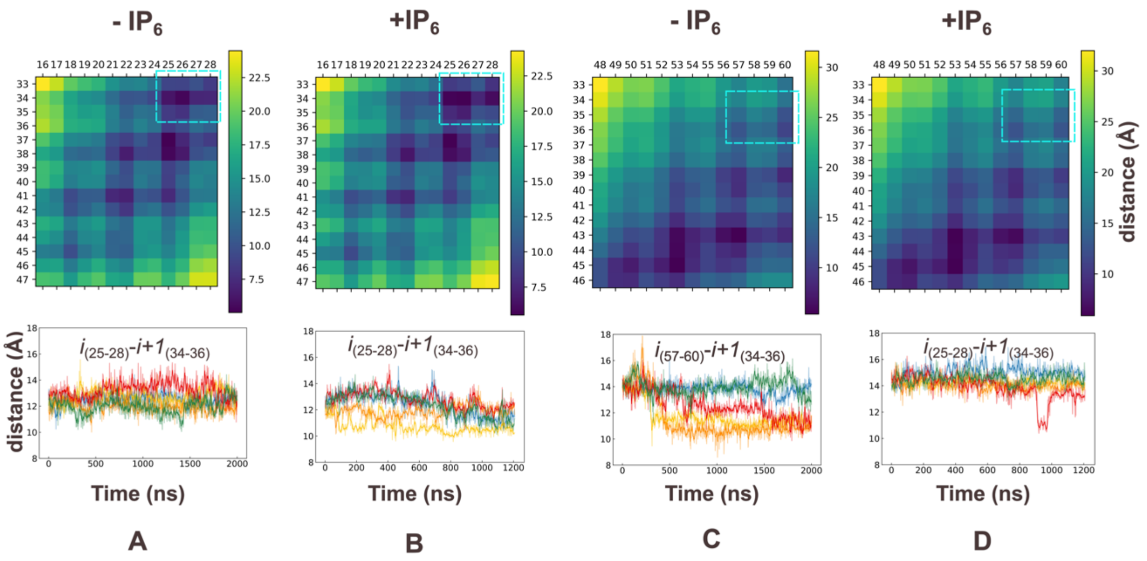


**Fig. S13. Comparison of hydrophobic core of the Pentamer NTD. (A)** Color map of contacts between helix 1 and neighboring helix 2 (top) and time series of distance between residues 25-28 of helix 1 and 34-36 of adjacent helix 2 (bottom). **(B)** Contact map for (+IP$_6$) simulations revealed interaction between the base of helix 1 and helix 2 (top), time course of residues 25-28 of helix 1 and 34-36 of adjacent helix 2 (bottom). **(C)** the base of CA helix 3 moves toward helix 2 of adjacent NTD adopting a hexamer-like state in the absence of IP$_6$ (top), time series analysis shows helix 3 of $CA_i$ contacting helix 2 of $CA_{i+1}$(bottom). **(D)** Pentamer NTD-NTD interface is preserved in (+IP$_6$)-simulations.

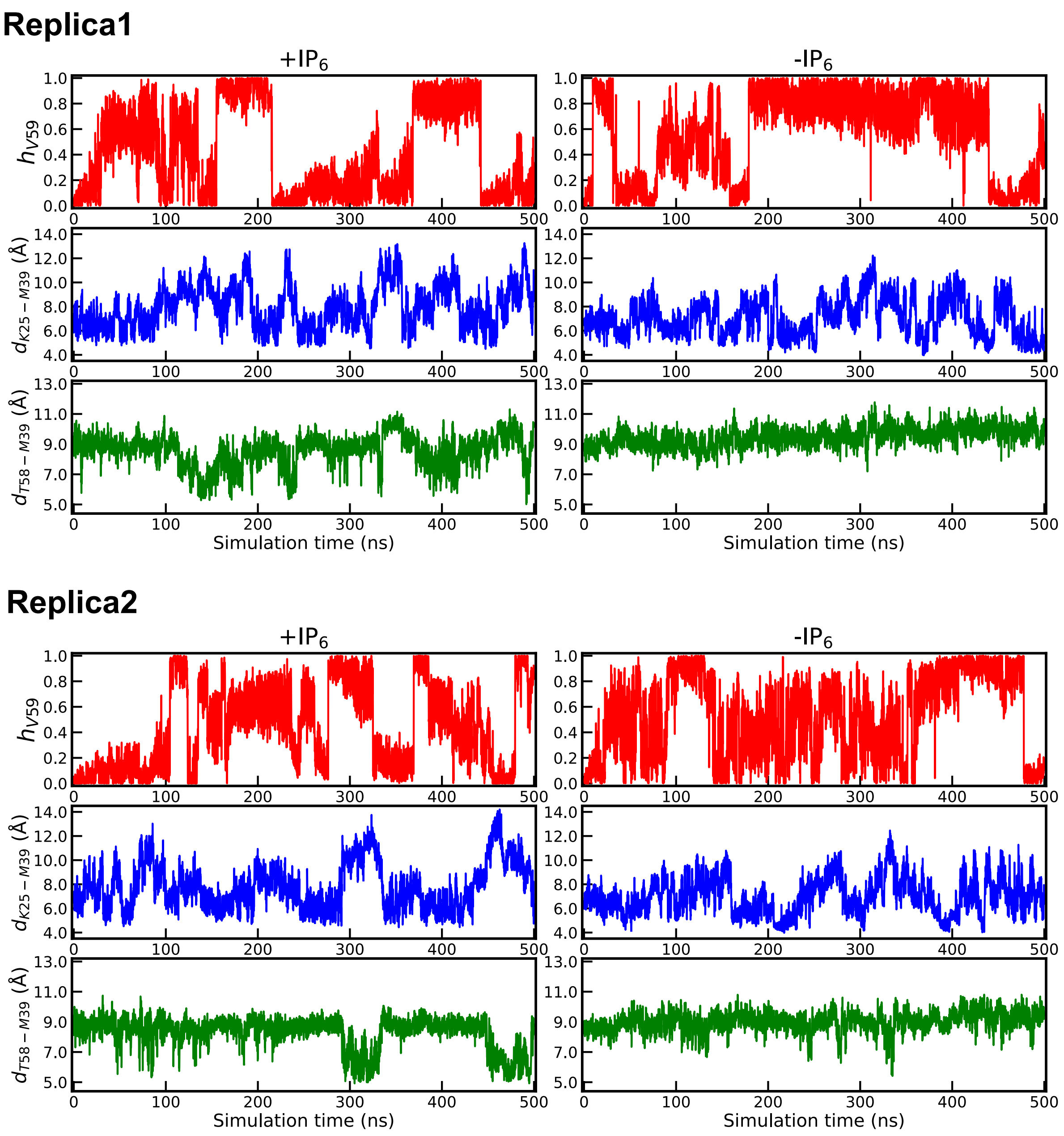


**Fig. S14.** Time series of the WT-MetaD simulation in the presence (+$IP_6$) and absence (-$IP_6$) of $IP_6$ bound to the pore. The time series of the helicity parameter ($h_{V59}$), the distance between K25 and M39 ($d_{K25-M39}$) of $CA_1$ and $CA_2$, and the distance between K58 and M39 ($d_{T58-M39}$) of $CA_1$ and $CA_2$, are shown in upper, middle, and lower panels, respectively.

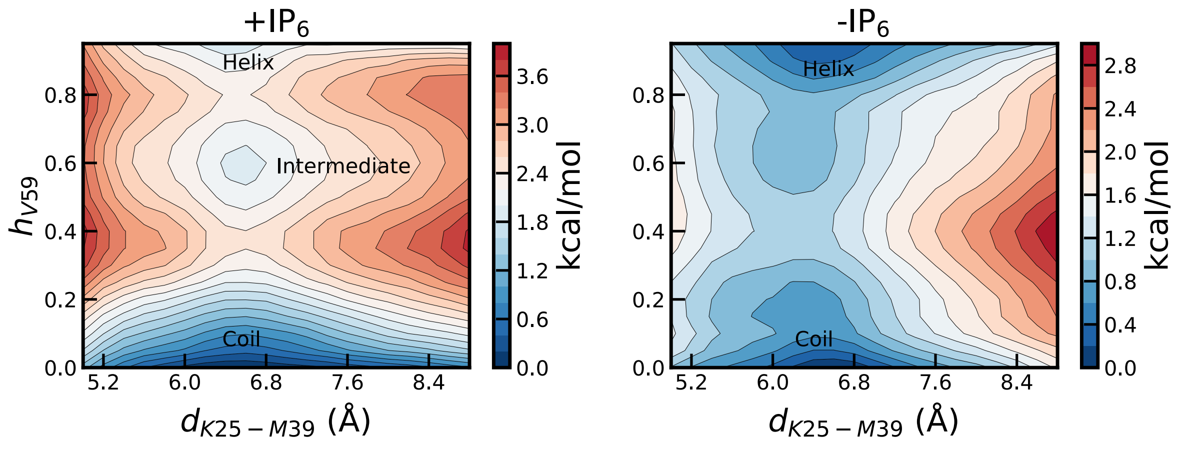


**Fig. S15.** 2-dimensional plot (Replica 2) of the PMF of the conformation landscape of the TVGG motif as a function of the helicity ($h_{V59}$) of V59, and distance between K25 of $CA_1$ and M39 of $CA_2$ denoted ($d_{K25-M39}$). The bar on the right has the color code for energy level contours in kcal/mole. The PMFs were calculated from the WT-MetaD simulations performed in the presence (+IP$_6$) and absence of (-IP$_6$). The PMFs of Replica 1 is shown in the main manuscript. The energy scale is 0.2 kcal/mol.

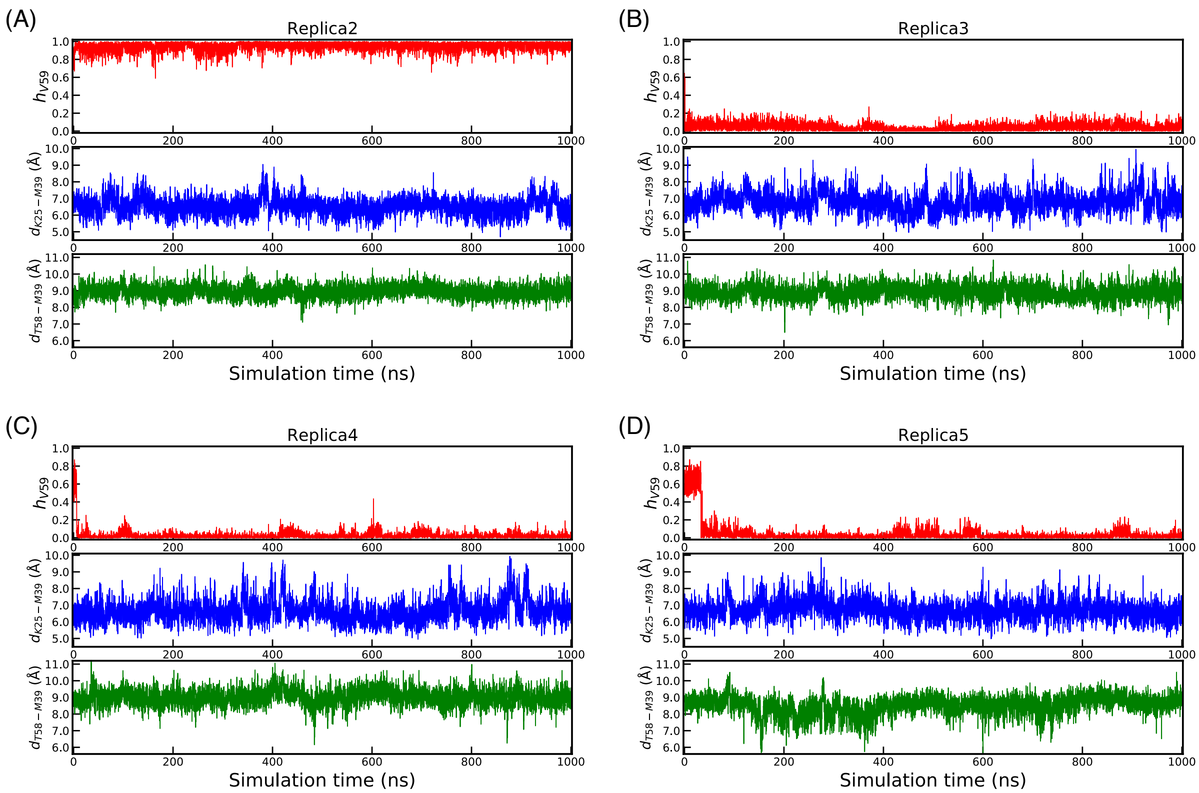

**Fig. S16. Dynamics of coil-to-helix transition from the intermediate partially folded state. (A-D)** Time evolution of the helicity parameter ($h_{V59}$), the distance between K25 and M39 ($d_{K25-M39}$) of $CA_1$ and $CA_2$, and the distance between K58 and M39 ($d_{T58-M39}$) of $CA_1$ and $CA_2$. The initial configuration corresponds to the intermediate partially folded state obtained from the WT-MetaD simulations in the presence of IP6. In the initial configuration of Replica2, the values of $h_{V59}$, $d_{K25-M39}$, and $d_{T58-M39}$ are 0.68, 6.5 Å, and 8.4 Å, respectively. In the initial configuration of Replica3, the values of $h_{V59}$, $d_{K25-M39}$, and $d_{T58-M39}$ are 0.64, 6.5 Å, and 8.8 Å, respectively. In the initial configuration of Replica4, the values of $h_{V59}$, $d_{K25-M39}$, and $d_{T58-M39}$ are 0.55, 6.1 Å, and 8.9 Å, respectively. In the initial configuration of Replica5, the values of $h_{V59}$, $d_{K25-M39}$, and $d_{T58-M39}$ are 0.58, 6.5 Å, and 8.7 Å, respectively.

**Movie S1:** Dynamics of the central pore of the capsid pentamer in the absence of $IP_6$ ($-IP_6$). The pentameric central pore contains R18 and K25 rings, each made of five electropositive residues in proximity. The pore radius is smaller for the R18 ring compared to the K25 ring. In both rings, the sidechains fluctuate rapidly to decrease the repulsive force within the ring by orienting the positive charges in opposite directions. This increases the fluctuation within the central ring.

**Movie S2:** Dynamics of the central pore of the capsid pentamer in the presence of $IP_6$ ($+IP_6$). In $IP_6$-bound pentameric, $IP_6$ coordinates the positive charges in the R18 and K25 rings to stabilize the central pore. Thus, the sidechain fluctuations in both rings are neutralized, bringing overall order to the pore. Electropositive sidechains in both rings point toward the center of the ring.
